# Therapeutic DNA Vaccine Targeting *Mycobacterium tuberculosis* Persisters Shortens Curative Tuberculosis Treatment

**DOI:** 10.1101/2024.09.03.611055

**Authors:** Styliani Karanika, Tianyin Wang, Addis Yilma, Jennie Ruelas Castillo, James T. Gordy, Hannah Bailey, Darla Quijada, Kaitlyn Fessler, Rokeya Tasneen, Elisa M. Rouse Salcido, Harley T. Harris, Rowan E. Bates, Heemee Ton, Jacob Meza, Yangchen Li, Alannah D. Taylor, Jean J. Zheng, Jiaqi Zhang, J David Peske, Theodoros Karantanos, Amanda R. Maxwell, Eric Nuermberger, Richard B. Markham, Petros C. Karakousis

## Abstract

*Mycobacterium tuberculosis (*Mtb) is one of the leading infectious causes of death worldwide. There is no available licensed therapeutic vaccine that shortens active tuberculosis (TB) disease drug treatment and prevents relapse, despite the World Health Organization’s calls. Here, we show that an intranasal DNA vaccine containing a fusion of the stringent response *rel*_Mtb_ gene with the gene encoding the immature dendritic cell-targeting chemokine, MIP-3α/CCL20, shortens the duration of curative TB treatment in immunocompetent mice. Compared to the first-line regimen for drug-susceptible TB alone, our novel adjunctive vaccine induced greater Rel_Mtb_-specific T-cell responses associated with optimal TB control in spleen, blood, lungs, mediastinal lymph nodes, and bronchoalveolar lavage (BAL) fluid. These responses were sustained, if not augmented, over time. It also triggered more effective dendritic cell recruitment, activation, and colocalization with T cells, implying enhanced crosstalk between innate and adaptive immunity. Moreover, it potentiated a 6-month TB drug-resistant regimen, rendering it effective across treatment regimens, and also showed promising results in CD4+ knockout mice, perhaps due to enhanced Rel-specific CD8+ T-cell responses. Notably, our novel fusion vaccine was also immunogenic in nonhuman primates, the gold standard animal model for TB vaccine studies, eliciting antigen-specific T-cell responses in blood and BAL fluid analogous to those observed in protected mice. Our findings have critical implications for therapeutic TB vaccine clinical development in immunocompetent and immunocompromised populations and may serve as a model for defining immunological correlates of therapeutic vaccine-induced protection.

**One sentence summary:** A TB vaccine shortens curative drug treatment in mice by eliciting strong TB-protective immune responses and induces similar responses in macaques.

## Introduction

Ten million people fall ill with tuberculosis (TB) every year (*1*). It remains among the leading single infectious causes of death, leading to over a million deaths annually, even though TB is curable (*1*). People living with HIV/AIDS (PLWHA) are ∼18 times more likely to develop active TB disease than people without HIV. The former group is at increased risk of adverse TB treatment outcomes, including treatment failure, recurrence, emergence of drug resistance, and death (*1*). The current six-month regimen, consisting of rifampin (R), isoniazid (H), pyrazinamide (Z), and ethambutol (E), has high efficacy against drug-sensitive TB. However, its length and complexity contribute to treatment interruptions that jeopardize cure and promote drug resistance (*2, 3*). Drug-resistant (DR) TB remains a public health crisis (*1*). Although newer TB drugs, such as bedaquiline (B), pretomanid (Pa), and linezolid (L), offer hope for improved treatment of DR-TB cases, strains resistant to each of these new drugs have already emerged, sounding a note of caution against reliance on currently available antibiotic therapy to cure DR-TB in the future (*4–6*).

Given all the above, the World Health Organization strongly encourages the development of therapeutic TB vaccines as a potential strategy to address TB treatment challenges better, setting as one of the priorities the shortening of drug treatment without subsequent relapse in patients with active TB disease (*7*). Few therapeutic TB vaccines, such as RUTI (*8*), *M. indicus pranii* (*9*), *M. vaccae* (*10*), ID93+GLA-SE (*11*), and *MIP-3*α*/rel_Mtb_*(*12*), have been studied for this purpose in preclinical models, showing a range of adjunctive therapeutic efficacy when combined with TB drugs. However, none of these vaccines has been shown yet to shorten the duration of curative TB treatment without subsequent relapse after treatment discontinuation.

We have previously shown that intranasal (IN) delivery of the therapeutic DNA TB vaccine targeting the *Mycobacterium tuberculosis* (Mtb) stringent response factor Rel_Mtb_ to dendritic cells (DCs), *MIP-3*α*/rel_Mtb,_* when combined with H, substantially decreased the lung bacillary burden compared to H alone (*12*). Here, we present that the adjunctive *MIP-3*α*/rel_Mtb_* vaccine led to lung culture-negativity more rapidly than RHZE alone, yielding a stable cure after treatment discontinuation and, in parallel, was more effective in reducing the lung bacillary burden when combined with BPaL than BPaL alone in Mtb-infected immunocompetent mice. We also observed similar adjunctive therapeutic efficacy in a CD4 knockout (KO) mouse model. IN immunization with the *MIP-3*α*/rel_Mtb_* vaccine enhanced DC recruitment and activation, which was accompanied by increased production of antigen-specific-T-cell-produced cytokines associated with optimal TB control. Similarly, nonhuman primates immunized with *MIP-3*α*/rel_Mtb_* vaccine exhibited an increase in antigen-specific TB protective cytokines, suggesting that this vaccination strategy may be a promising adjunctive tool for TB treatment shortening.

## Results

### Therapeutic IN *MIP-3*α*/rel_Mtb_* fusion vaccine potentiated the first-line drug-sensitive TB regimen in immunocompetent mice

In the current study, we aimed to test the efficacy of the IN *MIP-3*α*/rel_Mtb_* fusion therapeutic TB vaccine in combination with the complete first-line drug-sensitive TB regimen RHZE in C57BL/6 mice. Four weeks after Mtb aerosol infection, mice were treated daily with human-equivalent doses of oral RHZE for 6 weeks (Fig. 1A). Separate groups of mice received either the *MIP-3*α*/rel_Mtb_* fusion vaccine or the original vaccine expressing only the antigen (*rel_Mtb_*), and each was administered once weekly via the IN vs. intramuscular (IM) route for 3 weeks. As controls, apart from receiving mock treatment and vaccination regimens (“Control”), another group received RHZE only. An additional control group received a fusion vaccine containing the *MIP-3*α gene fused to the gene encoding Early Secreted Antigenic Target of 6 kDa protein (*MIP-3*α*/ESAT-6)*, an Mtb secretory protein and immunodominant and potent T-cell antigen, as opposed to Rel_Mtb,_ a stringent response antigen (*13–15*).

**Fig. 1.**
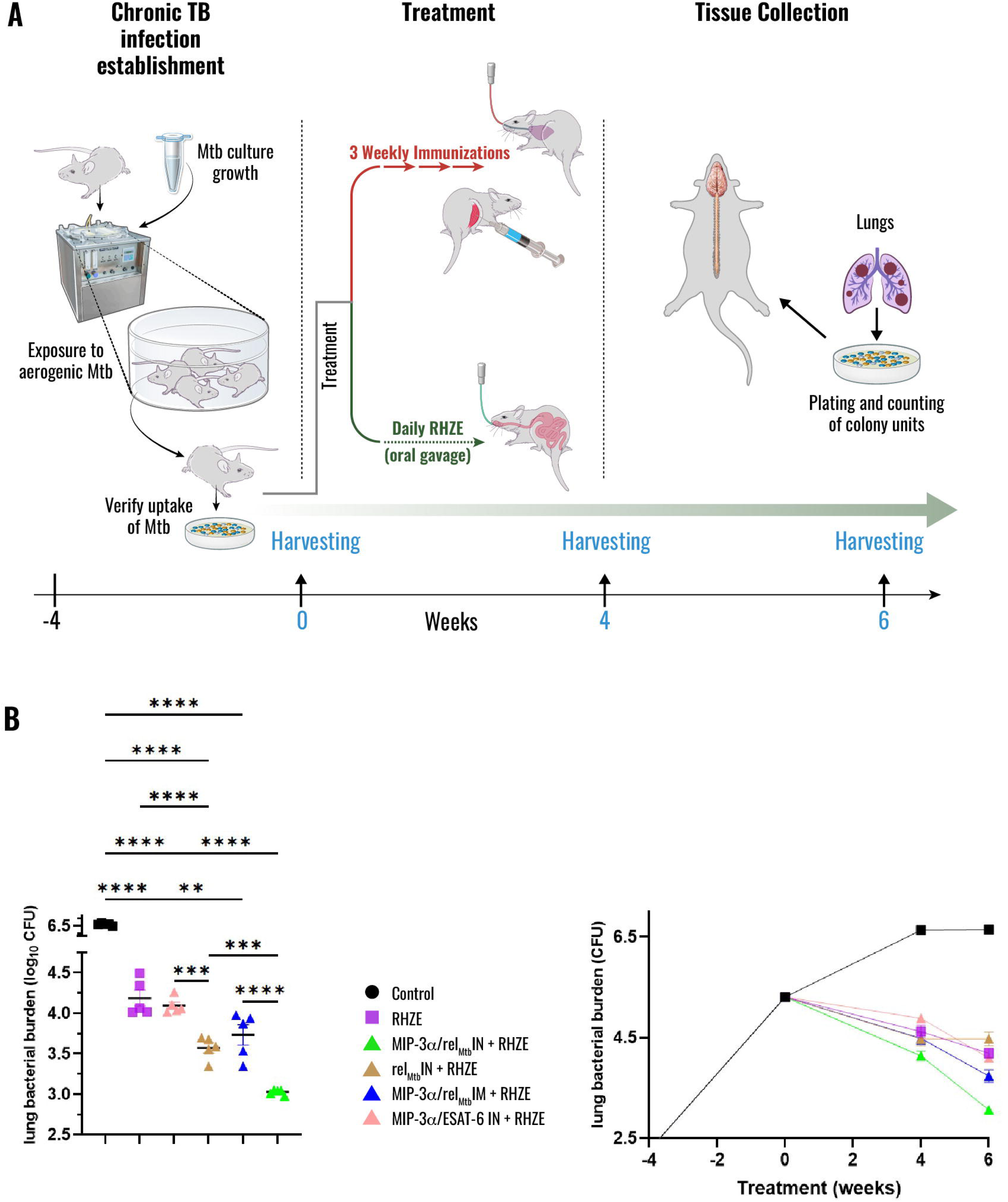
Therapeutic *IN MIP-3*α*/rel_Mtb_* fusion vaccine potentiated the first-line regimen for drug-susceptible TB in immunocompetent mice. (A) Timeline of *Mtb* challenge study; **(B)** Scatterplot; and **(C)** Timeline of lung mycobacterial burden at 4 and 6 weeks after the primary vaccination per vaccination group. Mtb, *Mycobacterium tuberculosis*, IM: intramuscular; IN: intranasal; ESAT-6, Early Secreted Antigenic Target; CFU, colony-forming units; RHZE, Rifampin-Isoniazid-Pyrazinamide-Ethambutol; **P<0.01, ***P<0.001, ****P<0.000.

At 6 weeks after primary vaccination, the most significant reduction in the lung mycobacterial burden was observed in the group receiving the IN *MIP-3*α*/rel_Mtb_*fusion vaccine along with RHZE compared to any other vaccination approach, RHZE alone or control (absolute reduction of mycobacterial burden: >0.5, 1.3 and 3.6 log_10_ CFU with P<0.001, P<0.0001 and P<0.0001, respectively, Figs 1B, C). Interestingly, no adjunctive therapeutic effect was observed after IN vaccination with *MIP-3*α*/ESAT-6* compared to RHZE alone (Fig. 1B). Thus, collectively, the IN *MIP-3*α*/rel_Mtb_* fusion vaccine was found to be the most effective vaccination strategy, resulting in the most significant mycobacterial reduction when combined with the first-line treatment for drug-sensitive TB.

### Adjunctive IN *MIP-3*α*/rel_Mtb_* fusion vaccine shortens the curative TB treatment in immunocompetent mice

Next, we investigated if the IN *MIP-3*α*/rel_Mtb_* fusion vaccine can shorten curative TB treatment in immunocompetent mice. Four weeks after Mtb aerosol infection, 10-20 male and female C57BL/6 mice per group were treated daily with human-equivalent doses of oral RHZE for 12 weeks (Fig. 2A). At 10 weeks after primary vaccination, the most significant reduction in the lung mycobacterial burden was observed to occur more rapidly in mice receiving the IN *MIP-3*α*/rel_Mtb_* fusion vaccine with RHZE compared to those receiving RHZE alone (additional reduction in mycobacterial burden: 1.98 log_10_ CFU, P<0.0001; Fig. 2B). At 12 weeks after primary vaccination, the lungs of all mice receiving the IN *MIP-3*α*/rel_Mtb_* fusion vaccine in combination with RHZE were culture-negative in contrast to those receiving RHZE group alone (0 log_10_ CFU vs. 2.58 log_10_ CFU, P<0.0001; Fig. 2B). RHZE was discontinued for 12 weeks upon completion of treatment to assess microbiological relapse in the lungs and spleens. None of the mice receiving the IN *MIP-3*α*/rel_Mtb_*fusion vaccine relapsed after the 12-week treatment discontinuation (0/20) in contrast to the RHZE group alone, in which all mice relapsed (20/20) (Figs 2C and 2D). Thus, IN delivery of the *MIP-3*α*/rel_Mtb_* fusion vaccine potentiates the activity of RHZE and shortens the duration of curative treatment with the first-line regimen in Mtb-infected mice.

**Fig. 2.**
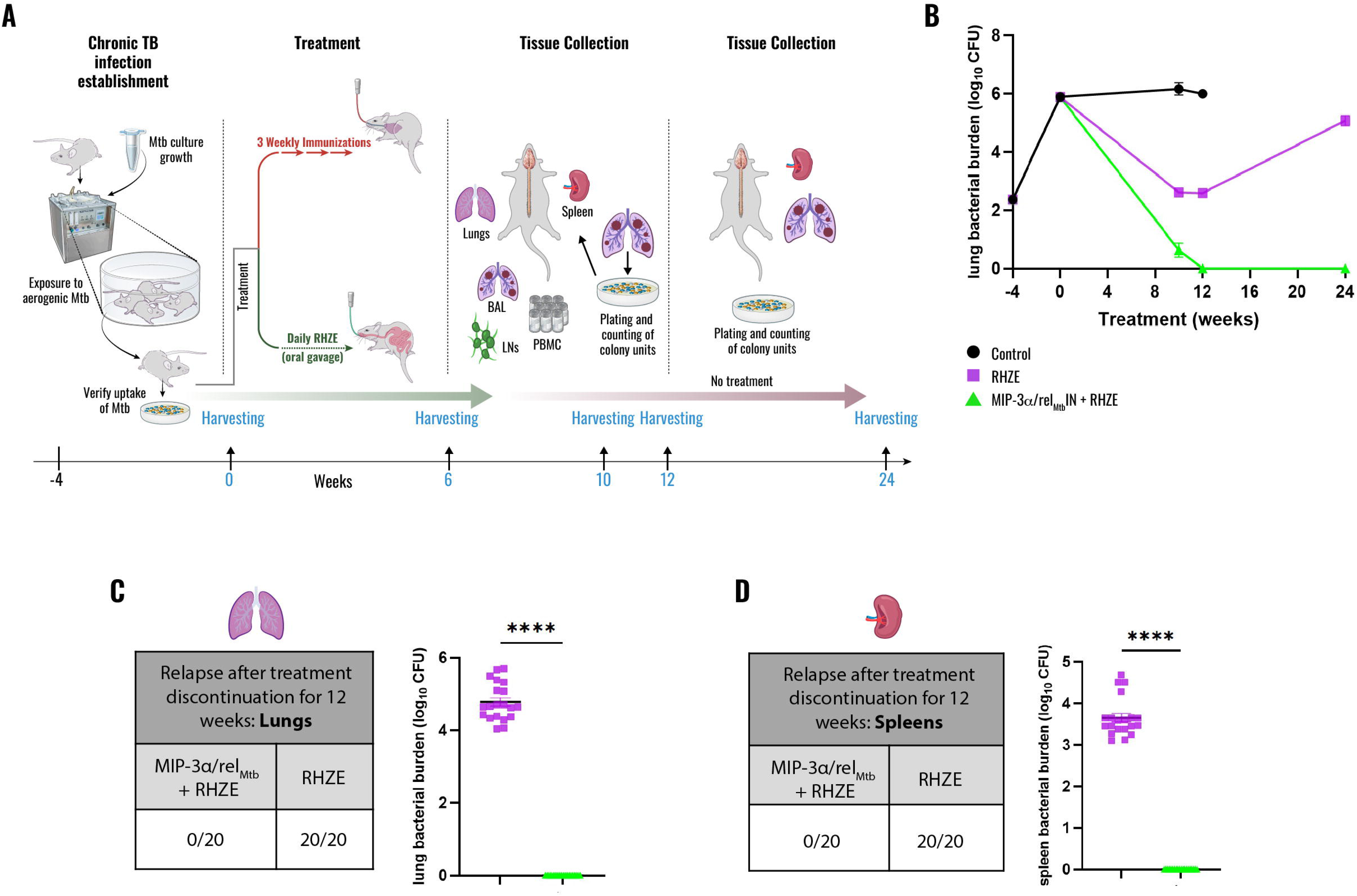
Adjunctive IN *MIP-3*α*/rel_Mtb_* fusion vaccine led to lung-culture-negativity more rapidly than RHZE alone, yielding a stable cure after 12 weeks of treatment in immunocompetent mice. (A) Timeline of Mtb challenge study; **(B)** Timeline of lung mycobacterial burden at 10, 12, and 24 weeks after the primary vaccination per vaccination group; **(C)** Scatterplot of the lung; and **(D)** spleen mycobacterial burden 24 weeks after primary vaccination (and 12 weeks after treatment discontinuation). Mtb, *Mycobacterium tuberculosi*s; IN: intranasal; CFU, colony-forming units; RHZE, Rifampin-Isoniazid-Pyrazinamide-Ethambutol; ****P<0.0001.

### Therapeutic IN *MIP-3*α*/rel_Mtb_* fusion immunization induced sustained TB-protective immune responses when combined with the first-line drug-sensitive TB regimen

We measured the immune responses across the different groups over time (6 and 12 weeks post-treatment; Fig. 3). Compared to the RHZE group alone, immunization with IN *MIP-3*α*/rel_Mtb_ f*usion vaccine induced significantly more Rel_Mtb_-specific CD4+ and CD8+ T cells producing IFN-γ, TNF-α, and IL17-A, which are important cytokines for optimal TB control (*16–18*), both systemically (spleen and peripheral blood mononuclear cells (PBMCs)) and at the site of infection (lungs, mediastinal lymph nodes (LNs) and bronchoalveolar lavage (BAL)) (Fig. 3). This vaccination strategy also resulted in increased percentages of total classical DCs, activated DCs, and their major subtypes (Classical DC type I and II) relative to RHZE alone, consistent with increased DC recruitment and activation, as well as enhanced antigen presentation and CD4+ and CD8+ T-cell response priming. These responses were augmented at 12 weeks post-treatment (Fig. 3).

**Fig. 3.**
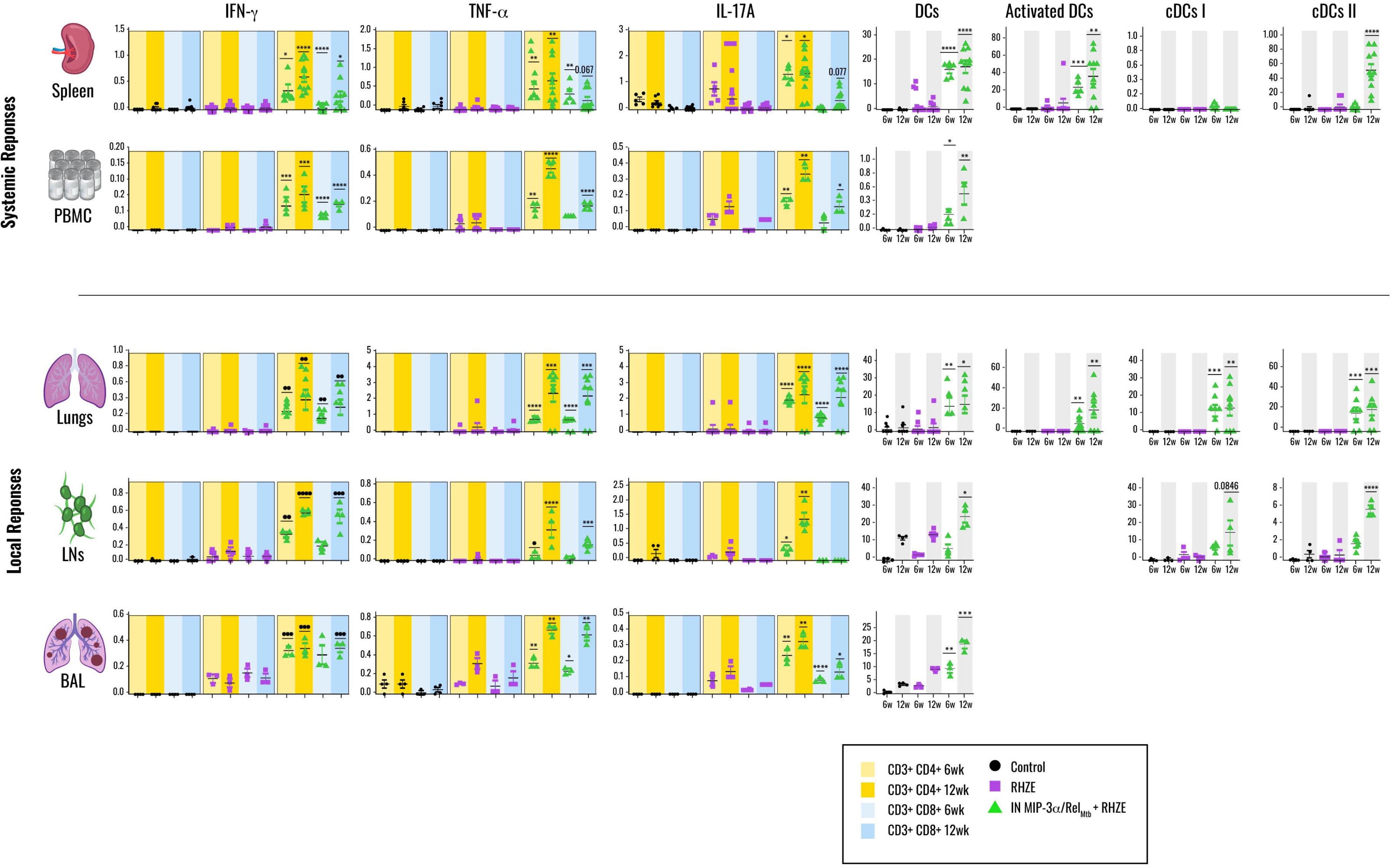
Therapeutic IN *MIP-3*α*/rel_Mtb_* fusion immunization induced sustained local and systemic TB-protective immune responses when combined with the first-line regimen for drug-susceptible TB regimen (RHZE) in immunocompetent mice. Six and 12-week Rel_Mtb_-specific CD4+ and CD8+ T-cell responses (IFN-γ, TNF-α, IL-17A) and percentage of DCs, activated DCs, cDCs I and II in spleen and PBMC (representing systemic responses) and lungs, LNs and BAL (representing local responses). Cytokines and DCs are expressed in percentages. IN, intranasal; RHZE, Rifampin-Isoniazid-Pyrazinamide-Ethambutol; DCs, total classical dendritic cells; cDCs I, classical dendritic cells type I; cDCs II, classical dendritic cell type 2; PBMC, peripheral blood mononuclear cells; LNs, mediastinal lymph nodes; BAL, bronchoalveolar lavage; 6w, 6 weeks; 12w, 12 weeks; *P<0.05; **P<0.01; ***P<0.001; ****P<0.0001.

### Immune profiles associated with mycobacterial control

Several simple linear regressions were used to test whether IN fusion vaccine-induced peak antigen-responsive T-cell-producing cytokines and/or dendritic cell proportions in various tissues (spleen, lungs, LNs, PBMC, and BAL) were associated with lung mycobacterial burden. Higher percentages of Rel_Mtb_-specific IL-17A+, IFN-γ+, and TNF-α+-producing T cells in PBMCs and BALs, along with Rel_Mtb_-specific IFN-γ+ and TNF-α+-producing T cells in LN and total classical DCs in BAL were highly and significantly associated with lower lung mycobacterial burden (Fig. S1). These findings suggest that these vaccine-induced immune responses may serve as indirect surrogates of optimal TB control.

### Therapeutic IN *MIP-3*α*/rel_Mtb_* fusion immunization promotes DC infiltration in the lungs and enhances their colocalization with T cells

We used multiplex immunofluorescence to quantify DCs and T cells in murine lungs collected 12 weeks after prime vaccination +/-RHZE treatment (Figs. 4 A, B). We also aimed to assess DC/T-cell colocalization as an indirect measure of possible enhanced DC/T-cell interactions (Figs 4A, B). Mice receiving the fusion vaccine and RHZE had a significantly higher percentage of DCs (defined as DCs/DAPI+ cells) in the lungs than those receiving RHZE alone (P=0.043, Fig. 4C), suggesting significantly higher local DC infiltration. The percentage of total (including non-antigen-specific) T cells did not differ significantly across the two groups, although a trend was observed, despite the small sample size in each group (P=0.352, Fig. S2, defined as T cells /DAPI+ cells). Next, we measured the numbers of DCs co-localized within 10 μm of T cells, a distance equal to a cell diameter, as a surrogate marker for cell-to-cell interactions. We found that the mouse group receiving the fusion vaccine and RHZE had a significantly higher number of DCs within 10 μm of a T cell than the group receiving RHZE alone (median number 44 DCs vs 17 DCs, P=0.019; Fig. 4D). These findings suggest that the IN fusion vaccine strategy induces local DC infiltration and enhances DC engagement of T cells.

**Fig. 4.**
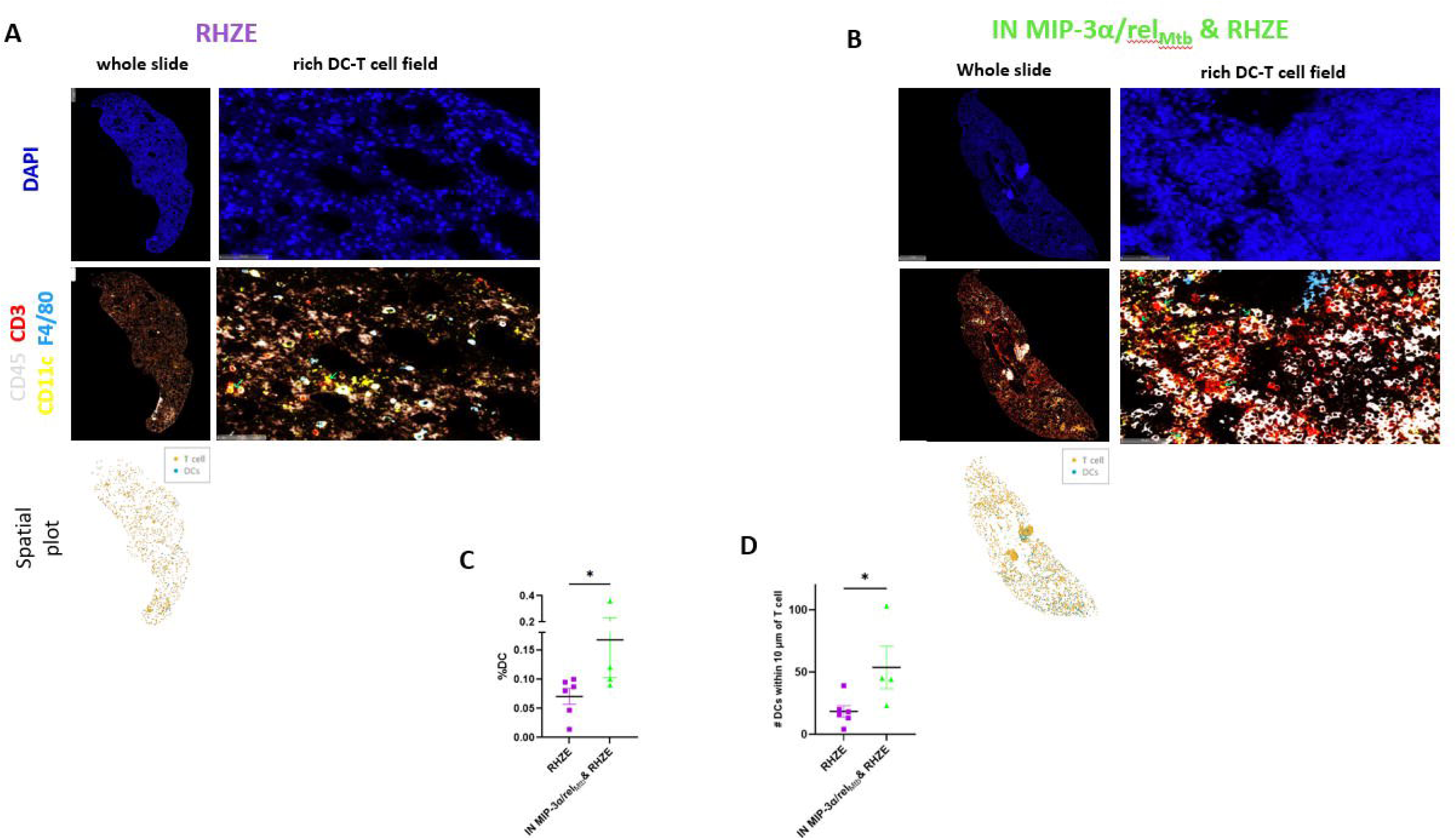
Therapeutic IN *MIP-3*α*/rel_Mtb_* fusion immunization induces local DC infiltration, enhancing colocalization with T cells in mouse lungs. A, B. Representative lung sections (whole slides, rich dendritic cell-T cell fields of view and DC-T cell spatial plots) from RHZE alone (A) *vs*. IN MIP-3α/rel_Mtb_ & RHZE after 12 weeks of treatment **(B)** stained with antibodies for DAPI+ only (dark blue, cell nuclei, top) or CD45+ (white, hematopoietic cells, bottom), CD45+CD3+ (red, T cells, bottom) and CD45+CD3-CD11c+F4/80-(yellow, dendritic cells, bottom). Green arrows indicate DC-T cell colocalization. **C, D.** Quantification of total dendritic cells per DAPI+ cells (%) **(C)** and **(D)** colocalization of dendritic-T cells defined as number of dendritic cells within 10 μm of T cells [RHZE alone (n=6) vs. IN MIP-3α/rel_Mtb_ & RHZE (n=4), Mann-Whitney test was run on medians, assessment was performed in whole slides]; DCs, dendritic cells; RHZE, Rifampin-Isoniazid-Pyrazinamide-Ethambutol; *P<0.05.

### Therapeutic IN *MIP-3*α*/rel_Mtb_* fusion vaccine potentiated the first-line drug-sensitive TB regimen in CD4 KO mice

To begin to understand the contribution of CD4+ T cells to the adjunctive therapeutic efficacy of the IN fusion vaccine, we next examined its antitubercular activity in combination with RHZE in a CD4 knockout (KO) mouse model, in particular because we noted robust classical DCs type 1 and CD8+ T-cell responses on top of DCs type 2 and CD4+T cell responses in most of the examined tissues in immunocompetent mice (Figs 3, 5). Four weeks after Mtb aerosol infection, each group of CD4 KO mice was treated daily with human-equivalent doses of oral RHZE for 6 weeks (Fig. 5A). The mouse group that received the IN fusion vaccine + RHZE had a significantly greater reduction in the lung mycobacterial burden compared to the group receiving RHZE alone after 6 weeks of treatment (CFU reduction difference: 0.75 log_10_, P<0.0001; Figs. 5B, C). A similar reduction in mycobacterial burden compared to RHZE alone was also observed in the spleens (CFU reduction difference: 1.56 log_10_, P<0.0001; Fig S3). These observations suggest that immunization with the IN fusion vaccine maintains its adjunctive therapeutic efficacy even in the setting of CD4 deficiency.

**Fig. 5.**
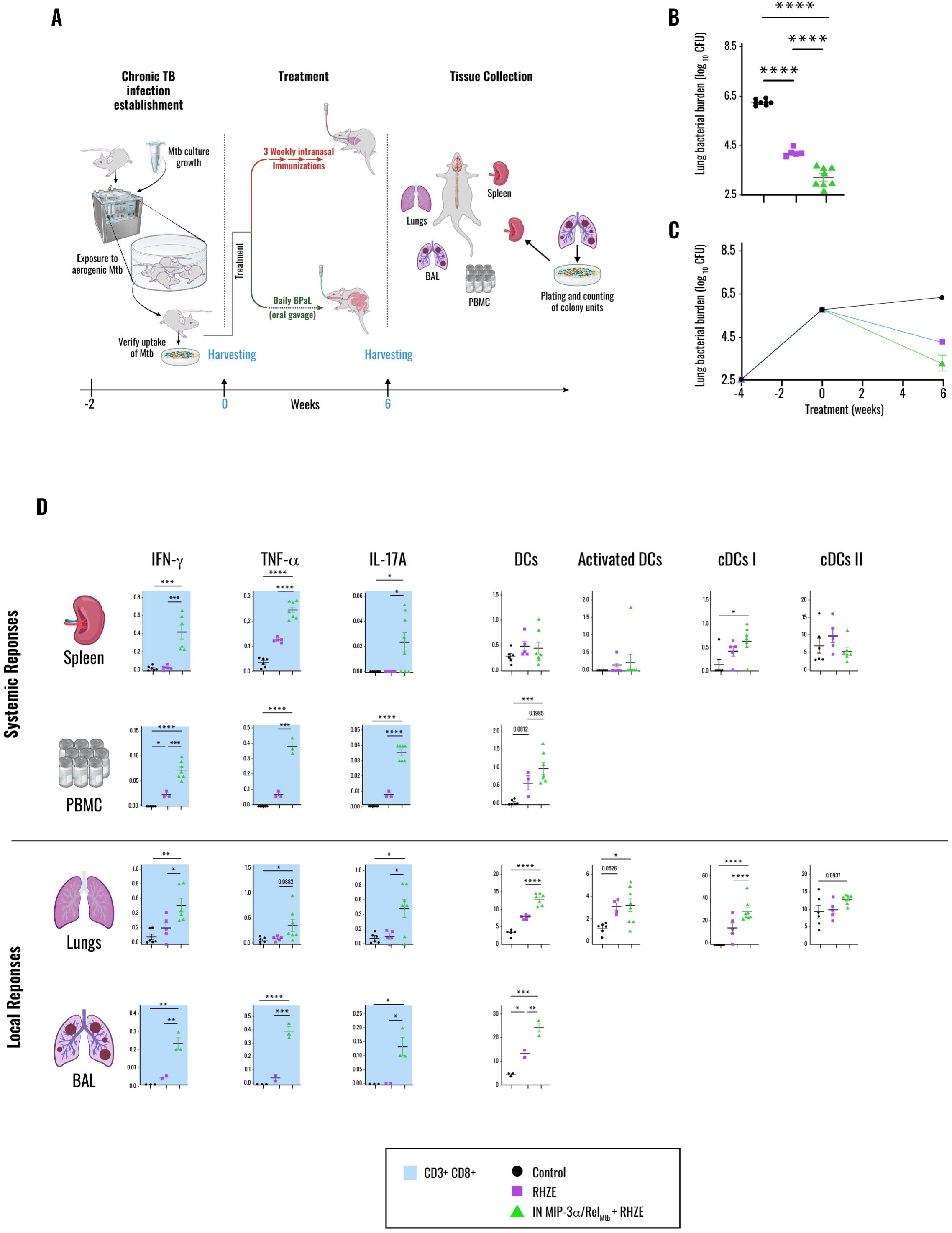
IN *MIP-3*α*/rel_Mtb_* DNA fusion vaccine potentiates the activity of RHZE and induces TB-protective immune responses in CD4 KO mice. (A) Timeline of Mtb challenge study; **(B)** Timeline of lung mycobacterial burden at -4, 0, and 6 weeks after treatment initiation; **(C)** Scatterplot of the lung mycobacterial burden at 6 weeks after treatment initiation; **(D)** 6-week rel_Mtb_-specific CD8+ T-cell responses (IFN-γ, TNF-α, IL-17A) and percentage of DCs, activated DCs, cDCs I and II in spleen and PBMC (representing systemic responses), lungs and BAL (representing local responses). Cytokines and DCs are expressed in percentages. Mtb, *Mycobacterium tuberculosi*s; IN: intranasal; CFU, colony-forming units; RHZE, Rifampin-Isoniazid-Pyrazinamide-Ethambutol; DCs, total classical dendritic cells; cDCs I, classical dendritic cells type I; cDCs II, classical dendritic cell type 2; PBMC, peripheral blood mononuclear cells; BAL, bronchoalveolar lavage; CD4 KO, CD4+ T cell knockout; *P<0.05; **P<0.01; ***P<0.001; ****P<0.0001.

### Therapeutic IN *MIP-3*α*/rel_Mtb_* fusion immunization induced TB-protective immune responses in the absence of CD4+ T cells

To gain insight into the therapeutic mechanism of action of the IN fusion vaccine in the CD4-deficient setting, we performed flow cytometry of samples derived from the lungs, BAL, spleens, and PBMCs. Similar to responses observed in immunocompetent mice, the IN fusion vaccine enhanced both systemic and local Rel_Mtb_-specific CD8+ T-cell responses (IFN-γ, TNF-α, IL17A) (Fig. 5D). It also led to increased numbers of classical DCs locally and in the periphery, increased local DC activation, and increased numbers of classical DC type 1 (Fig. 5D), which are well associated with cross-presentation and activation of CD8+ T cells, which in turn lyse host cells and kill *Mtb* directly, even in the absence of CD4+ T cells (*19–21*).

### Therapeutic IN *MIP-3*α*/rel_Mtb_* fusion vaccine potentiates a DR-TB regimen in immunocompetent mice

We next sought to determine if the adjunctive therapeutic efficacy of the IN *MIP-3*α*/rel_Mtb_*fusion vaccine is independent of antitubercular regimen and assessed its activity in combination with a novel, oral regimen for DR TB comprising BPaL (*22, 23*), compared to BPaL alone. Two weeks after Mtb aerosol infection, mice were treated daily with human-equivalent doses of oral BPaL for 6 weeks (Fig. 6A). We selected the subacute model of TB infection (i.e., higher Mtb inoculum with shorter incubation period before treatment initiation) instead of a chronic model to allow a sufficient dynamic range in which to assess any additional vaccine-induced mycobactericidal effect on top of the potent BPaL regimen. Relative to those receiving BPaL alone, mice receiving the IN fusion vaccine + BPaL had a significantly greater reduction in the lung mycobacterial load (CFU reduction difference: 0.7 log_10_, P<0.001; Figs. 6B, 5C) and a nonsignificant reduction in spleen mycobacterial burden (CFU reduction difference: 0.3 log_10_, P=0.8, Fig.S4) at 6 weeks of treatment. These findings suggest that IN immunization with the fusion vaccine offers adjunctive therapeutic efficacy independently of the drug treatment regimen.

**Fig. 6.**
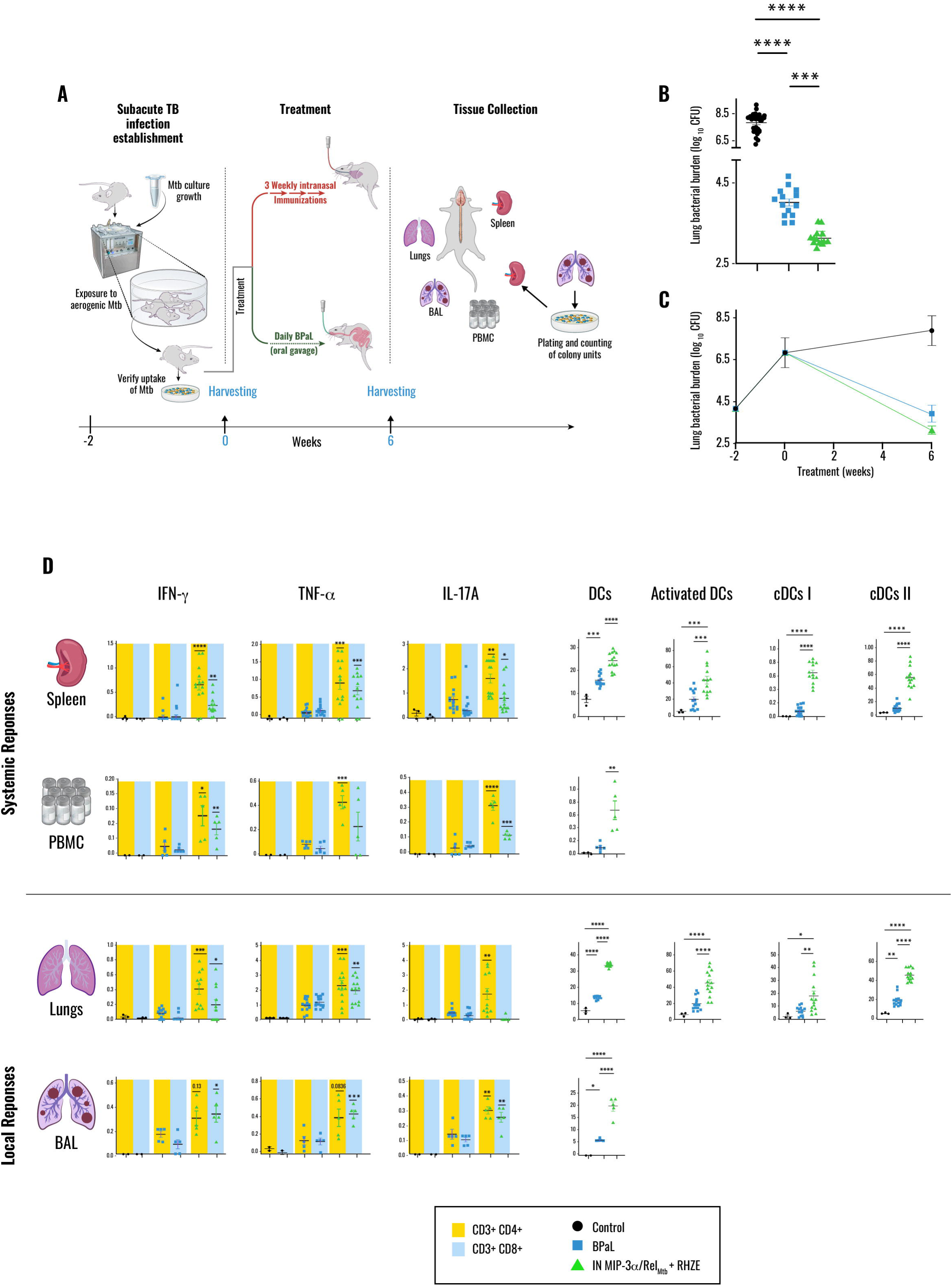
IN *MIP-3*α*/rel_Mtb_* DNA fusion vaccine potentiates the activity of the potent DR-TB regimen BPaL and induced TB-protective immune responses in immunocompetent mice. (A) Timeline of Mtb challenge study; **(B)** Timeline of lung mycobacterial burden at -2,0 and 6 weeks after treatment initiation; **(C)** Scatterplot of the lung mycobacterial burden at 6 weeks after treatment initiation; **(D)** 6-week rel_Mtb_-specific CD4+ and CD8+ T-cell responses (IFN-γ, TNF-α, IL-17A) and percentage of DCs, activated DCs, cDCs I and II in spleen and PBMC (representing systemic responses), lungs and BAL (representing local responses). Cytokines and DCs are expressed in percentages. Mtb, *Mycobacterium tuberculosi*s; IN: intranasal; CFU, colony-forming units; BPaL, Bedaquiline-Pretomanid-Linezolid; DC, total classical dendritic cells; cDCs I, classical dendritic cells type I; cDCs II, classical dendritic cell type 2; PBMC, peripheral blood mononuclear cells; BAL, Bronchoalveolar lavage; *P<0.05; **P<0.01; ***P<0.001; ****P<0.0001.

### Therapeutic IN *MIP-3*α*/rel_Mtb_* fusion immunization induces TB-protective immune responses when combined with a potent DR-TB regimen

Next, we measured both local and systemic immune responses following 6 weeks of treatment with BPaL +/- IN fusion vaccine (Fig. 6D). Similar to the RHZE study, compared to the BPaL-only group, IN fusion vaccine combined with BPaL led to increased percentages of Rel_Mtb_-specific CD4+ and CD8+ T cells producing IFN-γ, TNF- α, and IL17-A both systemically (spleen and PBMCs) and at the site of infection (lungs and BAL) (Fig. 6D). The IN fusion vaccine combined with BPaL also similarly resulted in increased numbers of total classical DCs, activated DCs, and their major subtypes (Classical DC type I and II) in contrast to BPaL alone (Fig. 6D).

### Testing the translational potential of the IN *MIP-3*α*/rel_Mtb_* fusion vaccine in nonhuman primates

To test the translational potential of this therapeutic vaccination strategy, we sought to investigate the immunogenicity of the IN *MIP-3*α*/rel_Mtb_* fusion vaccine in rhesus macaques, whose immune system closely resembles that of humans (*24–29*). We administered the vaccine in 3 doses, 3 weeks apart, at a dose of 1 mg IN per macaque, and measured the Rel*_Mtb_*-specific T-cell responses pre-vaccination and at 9 weeks and 24 weeks post-vaccination (3 and 18 weeks after the last vaccination, Fig. 7A). IN immunization with the *MIP-3*α*/rel_Mtb_* fusion vaccine led to a relative or statistically significant increase in Rel_Mtb_-specific IFN-γ-, TNF-α-, or IL17-A producing-CD4+ and CD8+ T-cell percentages at 9 weeks compared to pre-vaccination measurements in PBMCs or BALs (Fig.7B). All of these immune parameters were also found to be inversely correlated with the lung mycobacterial burden in our mouse TB challenge studies (Fig. S1). Thus, these findings suggest that the IN *MIP-3*α*/rel_Mtb_* fusion vaccine elicited antigen-specific responses associated with optimal TB control in nonhuman primates, the gold standard preclinical model for TB vaccines (*24–29*). Notably, those responses did not decline substantially over time, as measured at 24 weeks (Fig. S5).

**Fig. 7.**
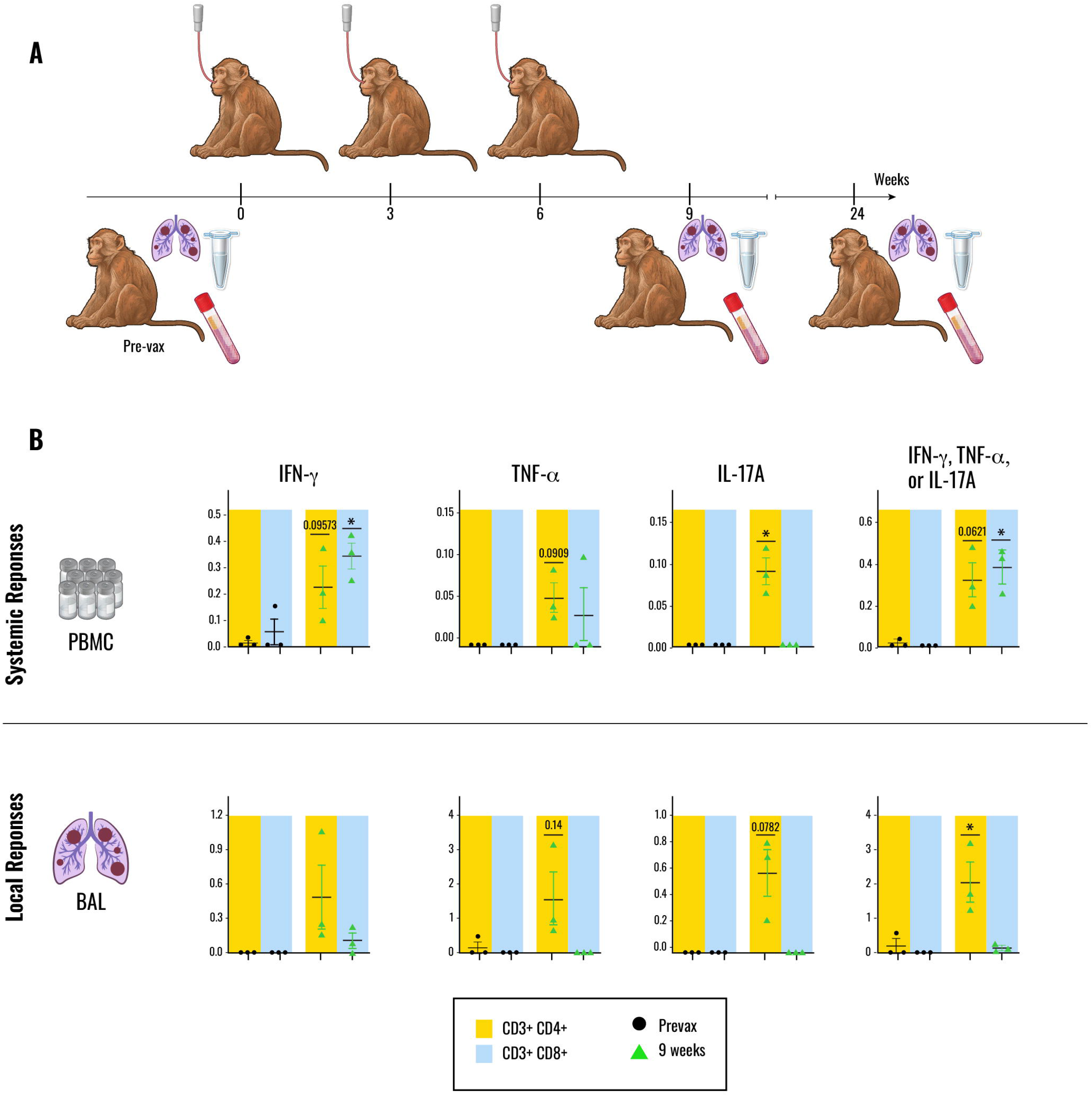
IN *MIP-3*α*/rel_Mtb_* DNA fusion vaccine induced TB-protective immune responses in nonhuman primates. **(A)** Timeline of the immunogenicity study; **(B)** 9-week Rel_Mtb-_specific CD4+ and CD8+T-cell responses (IFN-γ, TNF-α, IL-17A) in PBMC and BAL. Cytokines are expressed in percentages. IN, intranasal; PBMC, peripheral blood mononuclear cells; BAL, bronchoalveolar lavage; prevax, prevaccination; *P<0.05.

## Discussion

In this work, our preclinical data highlight that the IN *MIP-3*α*/rel_Mtb_* fusion vaccine is a promising adjunctive therapeutic tool to shorten the duration of treatment with the first-line regimen for drug-susceptible TB. With this vaccination strategy, local antigen-specific T-cells producing protective cytokines detected both locally and systemically were associated with this therapeutic benefit in our Mtb-infected mouse studies. This novel vaccination approach was also effective when combined with the DR-TB regimen, BPaL, and when administered in the complete absence of CD4+ T cells. Immunization of uninfected rhesus macaques, the closest experimental TB model to humans, induced similar sustained antigen-specific immune responses, suggesting that this vaccine may be a promising therapeutic tool in the clinical setting.

In the setting of TB disease, DCs are fewer and dysfunctional (*30–32*), leading to altered cellular responses. In our novel vaccination strategy, we have fused the chemokine MIP-3α/CCL20 with the Mtb antigen of interest, Rel_Mtb_ (*12*), to target it more efficiently to the DCs via the CCR6 receptor, in an effort to enhance the crucial crosstalk between innate and adaptive immune responses against Mtb, and to optimize cross-presentation to CD4+ and CD8+ T cells (*33*), an essential component of immunity for optimal Mtb control (*16, 34*). Acknowledging the foremost hurdle to develop not only an effective but also a durable therapeutic vaccine against pulmonary TB, we chose to administer this fusion vaccine via the IN route based on extensive literature showing that respiratory mucosal immunization induces long-lasting, local antigen-specific Mtb-protective T-cell responses in contrast to parenteral administration routes (*32, 35, 36*). Indeed, in the current study, IN immunization with our novel fusion vaccine shortened the duration of curative TB treatment, increased DC recruitment and activation, and elicited robust antigen-specific TB-protective T-cell responses systemically and locally compared to RHZE alone. Interestingly, these robust immune responses were sustained, if not augmented, over time.

HIV-induced CD4+ T-cell dysfunction and depletion significantly increases the risk of TB recurrence, even in the context of antiretroviral therapy (*37*). Compared with DCs from healthy donors, DCs derived from the peripheral blood of HIV-infected individuals at different stages of infection have significantly reduced efficiency in stimulating allogeneic T cells, likely due to decreased activation (*38*). It is also important to note that the stimulation of Mtb-specific CD8+ T cells *in vivo* and *in vitro* does not necessarily require CD4+ T-cell help and may be facilitated directly through DCs either through direct loading of MHC I molecules (*20*), or through cross-priming (*21*). CD8+ T cells specific for mycobacterial antigens can produce IFN-γ, lyse host cells, and attack Mtb directly (*20*). We found that our novel IN *MIP-3*α*/rel_Mtb_* fusion vaccine increased DC activation systemically and locally, eliciting robust CD8+T-cell responses in addition to potent CD4+ T-cell responses, and promoted lung infiltration of enhanced type I conventional DCs, a subset which is highly effective in activating cytotoxic lymphocytes, including CD8+ T cells (*39*). Given all of the above, we tested the efficacy of this vaccine in Mtb-infected CD4+ KO mice and found that it offered substantial adjunctive therapeutic benefit. Despite the absence of CD4+ T cells, robust antigen-specific CD8+ T-cell-producing TB-protective cytokines were elicited in the context of higher numbers of total cDCs driven primarily by cDCs I. Although promising, this vaccination strategy requires additional testing in preclinical models of TB/HIV co-infection.

BPaL is a potent DR-TB drug regimen that was recently approved by WHO in the context of the ZeNiX trial, demonstrating cure rates >90% when BPaL was administered for just 6 months with a lower, non-toxic dose of L(*22*). However, isolates resistant to B and Pa have been increasingly reported, highlighting the potential for spreading drug resistance in real-world implementation (*40, 41*). Thus, DR TB is another compelling setting to develop effective host-directed therapies against TB. This work shows that the novel IN *MIP-3*α*/rel_Mtb_*fusion vaccine combined with BPaL is more effective in reducing the mycobacterial burden than BPaL alone, as observed when the same vaccine is combined with RHZE, suggesting that it may also be effective in shortening curative treatment for DR-TB, since its activity is independent of drug resistance.

DNA vaccines have multiple advantages, including cost-efficient production, ease of manufacturing, safety in handling, and long shelf life, which are compelling features, especially for developing countries where TB is endemic (*42*). The challenge of their limited immunogenicity in nonhuman primates and humans has historically undermined their use, but recent developments in the context of the COVID-19 pandemic have shed more light on the scientific value of DNA vaccines, revealing several of these to be safe and immunogenic (*43–45*). Indeed, in this current work, apart from the therapeutic efficacy in mouse models, we found that our DNA IN fusion vaccine is immunogenic in nonhuman primates. One possible explanation of its effective delivery and immunogenicity is the fusion with the MIP-3α chemokine, which has been shown to be crucial in driving DC recruitment to the nasal mucosa (*46*).

Our study has some limitations. We did not investigate the therapeutic efficacy of our DNA fusion vaccine in a preclinical model with human-like TB pathology, such as the C3HeB/FeJ mouse model, which develops necrotic TB lung granulomas and even cavitary TB disease (*47*), the latter of which is a poor prognostic factor for TB treatment responses (*48*). These studies are currently ongoing. Additional macaque studies are also needed, including Mtb challenge studies, to confirm the therapeutic efficacy of the vaccine in the gold standard animal model before its evaluation in humans.

In conclusion, this study provides a paradigm shift towards developing safe therapeutic DNA TB vaccines focused on shortening TB drug treatment and decreasing relapse rates in various settings, including for both drug-susceptible and DR-TB without promoting drug resistance or adverse drug reactions in immunocompetent and immunocompromised hosts. The data also support the successful delivery of this DNA vaccine not only in small animal models but also in nonhuman primates, making it promising for successful clinical application. Finally, this study also provides a new framework for understanding the immunological correlates and mechanisms of TB control.

## Materials and Methods

### Ethics statement

All animal studies were performed per the protocols approved by the Johns Hopkins Animal Care and Use Committee of the Johns Hopkins School of Medicine. Macaques were housed and cared for following local, state, federal, and institute policies in facilities accredited by the American Association for Accreditation of Laboratory Animal Care (AAALAC) under standards established in the Animal Welfare Act and the Guide for the Care and Use of Laboratory Animals. Macaques were monitored for physical health, food consumption, body weight, and temperature. All experiments with Mtb in mice were conducted in Institutional Biosafety Committee-approved BSL3 and ABSL3 facilities at The Johns Hopkins University School of Medicine using recommended positive-pressure air respirators and protective equipment.

### Mouse Mtb infection studies

Four-to-six-week-old male and female C57BL6 mice and four-to-six-week-old female CD4 KO mice (homozygous for the Cd4tm1Mak targeted mutation) were purchased from The Jackson Laboratory. The mice were housed in individually ventilated cages maintained on a 12:12h light/dark cycle with free access to food and water. They were monitored at least weekly, recording their weights and assessing their general appearance. They were infected between 6 and 8 weeks of age via the aerosol route using the Glas-Col Inhalation Exposure System (Terre Haute, IN), and this procedure did not require anesthesia. A fresh aliquot of Wild-type *Mtb* H37Rv was used for each infection and diluted in 7H9 broth (Difco, Sparks, MD) with 10% oleic acid-albumin-dextrose-catalase (OADC), glycerol, and Tween-80 to achieve the desired inoculum per experiment. (*3*) The CFU implantation goal for the chronic TB infection model was ∼2 log_10_ of Mtb bacilli; for the subacute TB infection model, the goal was ∼ 4 log_10._ To reduce intergroup variability, mice assigned to the same experiment were infected together or evenly distributed between infection cycles and randomly assigned to each group per experiment. On the day after infection, at least 5 mice per experiment were sacrificed to determine the number of CFUs implanted into the lungs.

### Mouse treatments

After 14 (BPaL experiment) or 28 (RHZE experiments) days of infection, at least 5 animals per experiment were sacrificed to determine the CFU number present at the start of treatment. On the same day, for the RHZE experiments in C57BL6 and CD4 KO mice, the mice were randomized to receive human-equivalent doses of H (10 mg/kg), R (10 mg/kg), Z (150 mg/kg), and E (100 mg/kg) once daily dissolved in distilled water or distilled water only (control group) (*49*). The Z solution was gently heated in a 55°C water bath and vortexed to dissolve before treating mice. To minimize drug-drug interactions, R was administered at least 1h before Z. For the BPaL experiment in C57BL6 mice, the mice were randomized to receive B (25 mg/kg, formulated in 20% hydroxypropylb-cyclodextrin solution acidified with 1.5% 1N HCl) (*50*), Pa (50 mg/kg, prepared in the CM-2 formulation)(*51*), and L (50 mg/kg, prepared in 0.5% methylcellulose) (*50*) once daily or vehicle mix only (control group). The BPaL doses were selected to match the optimal human doses to minimize toxic effects while maintaining efficacy (*22, 52*). To minimize drug-drug interactions, L was administered 4h after BPa. All drugs in all experiments were administered as indicated, once daily, by orogastric gavage, in a total volume of 0.2-0.3 ml/mouse daily.

### Mouse vaccinations

At the time of drug treatment initiation, mice were randomized to receive the *rel_Mtb_* vaccine or the fusion *MIP-3*α*/rel_Mtb_* vaccine by the IM (10 μg in 50 μL in each quadriceps, followed by electroporation) or IN (100 μg in 50 μL in each nostril) route as described previously. (*12*) For the control or drug-only treatment groups, the mice received mock vaccination with IN PBS (50 ul per nostril per mouse). The mice were vaccinated three times at one-week intervals. IM or IN delivery of each plasmid followed adequate anesthesia of mice by vaporized isoflurane.

### Fluid/ Tissue Collection and bacterial enumeration

Mice were sacrificed 6, 10, 12, or 24 weeks after drug treatment and/or immunization series. For the efficacy time points/experiments, half spleens, right lower lungs, mediastinal LNs were harvested and processed into single-cell suspensions after mechanical disruption and filtration through a 70-μm cell strainer. Lung tissue was additionally digested using collagenase, Type I (ThermoFisher Scientific), and DNase (Sigma-Aldrich) for 20 min at 37 °C (*12*). Whole blood was collected through terminal cardiac puncture and processed into PBMC (*12*). BAL was performed, fluid was collected, and cellular component was isolated as a single-cell suspension (*53*). Single-cell suspensions were resuspended in warm R10 (RPMI 1640 with 2 mM-glutamine, 100 U ml−1 penicillin, 100 μg ml−1 streptomycin, and 10%heat-inactivated FBS; Atlantic Biologicals) for flow cytometry analysis as detailed below. The right upper lung per mouse was homogenized using glass homogenizers. Serial tenfold dilutions of lung homogenates in PBS were plated on 7H11 selective agar (BD) for colony-forming unit (CFU) enumeration. Plates were incubated at 37^0^C, and CFU were counted 4 weeks later by at least 2 investigators (*12*). L whole lung were fixed by immersion in 10% neutral buffered formalin for at least 48h, paraffin-embedded, sectioned, and underwent quadruple immunolabeling for DAPI+CD45+CD3+CD11c+F4/80+ as detailed below. For the relapse time point (24-week time point), whole spleens and whole lungs were plated for CFU enumeration.

### Macaque vaccinations, procedures, and sample collection

One-to-three-year-old three male or female rhesus macaques purchased from Johns Hopkins University Breeding Farm were used in the immunogenicity study (without Mtb infection). All macaques were housed in single-species harem breeding groups or same sex juvenile/young adult groups. Animal enclosures consisted of runs with concrete flooring or raised corncrib cages. All macaques had indoor and outdoor access, were fed a standard commercial diet (rhesus macaques: 5049 Fiber-Plus Monkey Diet, LabDiet) and rotating food enrichment items including fresh fruits, vegetables, and dried fruit treats. Animals were provided water ad libitum. Annual colony health screening included intradermal tuberculin testing and serology for Macacine herpesvirus 1 (B virus), simian immunodeficiency virus, simian T-cell leukemia virus, and simian retrovirus. All animals were consistently negative on tuberculosis testing and viral serology. Macaques were sedated with 10 mg/kg ketamine IM to receive IN immunization with the fusion *MIP-3*α*/rel_Mt_*_b_ vaccine. Animals were monitored closely throughout sedation until complete recovery. The vaccine was delivered as 500 μg plasmid DNA in 250 μl dripped into each nostril via pipette. Each macaque received three vaccinations administered at three-week intervals (week 0, week 3, week 6). Before prime immunization, at nine weeks and 24 weeks after the final vaccination, veterinary staff collected bronchoalveolar lavage (BAL) fluid and whole blood. BAL fluid was collected after ketamine and dexmedetomidine sedation and intubation. A single-use pediatric suction catheter (Airlife Tri-Flo Suction Catheter with Control Port, Carefusion, Yorba Linda, CA) was inserted through the endotracheal tube connector and blindly passed through the trachea and into a mainstem bronchi. The catheter was passed until resistance was felt, indicating the catheter had wedged into a distal bronchus. A 20 mL syringe containing the lavage fluid was attached to the end of the catheter and infused over 1 to 2 s. Immediately after the infusion, the aliquot was manually aspirated into the syringe using gentle pulsating suction. This process was repeated 2-3 times per animal. After the procedure, anesthesia was reversed with atipamazole hydrochloride (Antisedan, Zoetis, Kalamazoo, MI), and all animals were extubated and allowed to recover. Blood draws of 4 ml were obtained at the points of BAL (under sedation). BAL samples were processed into BAL single-cell suspensions by centrifugation and blood samples into PBMCs by Ficoll-paque PLUS gradient separation (GE Healthcare Biosciences) for flow cytometry analysis.

### Multiparameter flow cytometry

Single-cell suspensions from spleens, lungs, draining LNs, BAL, and PBMCs derived from mice or macaques were prepared as above. For LNs, BALs, and PBMCs, samples were pooled into 2-7 samples to achieve at least 10^5^ alive cells/sample. When applicable, each tissue was stimulated with purified recombinant Rel_Mtb_ protein at 37oC (3, 19) for various time intervals, from 12 hrs (IFN-γ, IL-17α, IL-2) to 24 hrs (ΤΝF-α), depending on the cytokine of interest (*12*). For Intracellular Cytokine Staining (ICS), GolgiPlug cocktail (BD Pharmingen, San Diego, CA) was added for an additional 4 hours after stimulation (total, 16 and 28 hours, respectively). Cells were collected using FACS buffer (PBS + 0.5% Bovine serum albumin (Sigma-Aldrich, St. Louis, MO), stained with Zombie NIR™ Fixable Viability Kit (Biolegend Cat. No.: 423106) for 30 min, and washed with PBS buffer. Surface proteins were stained for 20 min, the cells were fixed and permeabilized with buffers from Biolegend intracellular fixation/permeabilization set following manufacturer protocols (Cat. No. 421002), intracellular proteins were stained for 20 min, and samples were washed and resuspended with FACS buffer. The following anti-mouse mAbs were used: PE-conjugated anti-CD45 (Biolegend Cat. No. 103106), BV650-conjugated anti-CD45 (Biolegend Cat. No. 103151), PerCPCy5.5 conjugated anti-CD3 (Biolegend Cat. No. 100218), PE/Dazzle-conjugated anti-CD4 (Biolegend Cat. No. 100566), Alexa700 conjugated anti-CD8 (Biolegend Cat. No. 100730), PECy7-conjugated anti-TNF-α, (Biolegend Cat. No. 506324), APC-conjugated anti-IFN-γ, (Biolegend Cat. No. 505810), PE-conjugated anti-IL-17A (Biolegend Cat. No. 506904), BUV395-conjugated anti-IL-17A (BD Biosciences Cat. No. 565246), BUV563-conjugated anti-CD19 (BD Biosciences Cat. No. 749028), BUV496-conjugated anti-NK-1.1 (BD Biosciences Cat. No. 741062), BV605-conjugated anti-Ly-6G (Biolegend Cat. No. 127639), BV421-conjugated anti-CD11c (Biolegend Cat. No. 117330), BUV395-conjugated anti-CD24 (BD Biosciences Cat. No. 744471), BUV805-conjugated anti-CD11b (BD Biosciences Cat. No. 741934), BV785-conjugated anti-XCR1 (Biolegend Cat. No. 148225), BV650-conjugated anti-CD80 (Biolegend Cat. No. 104732), BV510-conjugated anti-MHCII (Biolegend Cat. No. 107636), FITC-conjugated anti-CD103 (Biolegend Cat. No. 121420). The following anti-non-human primates or human antibodies were used: FITC-conjugated anti-CD3 (ThermoFisher Scientific Cat. No. APS0308), Alexa700-conjugated anti-CD4 (ThermoFisher Scientific Cat. No. 56-0048-41), APC-anti-CD8 (ThermoFisher Scientific Cat. No. MA5-44089), PercPCy5.5-conjugated anti-TNF-α (ThermoFisher Scientific Cat. No. 45-7349-42), PE-Cy7-conjugated anti-IFN-γ (ThermoFisher Scientific Cat. No. 25-7319-82) and PE-conjugated anti-IL-17A (ThermoFisher Scientific Cat. No. 12-7178-42). The Attune™ NxT (Thermo Fisher Scientific, Waltham, MA), BD, BD Biosciences LSRII/Fortessa flow cytometer were used. Flow data were analyzed by FlowJo Software (FlowJo 10.8.1, LLC Ashland, OR). Gating strategies can be found in Figs S6 and S7.

### Immunofluorescence

Quadruple immunolabeling for CD45+CD3+CD11c+F4/80 was performed on formalin-fixed, paraffin-embedded sections derived from Mtb-mouse mouse lungs on a Ventana Discovery Ultra autostainer (Roche Diagnostics). Following dewaxing and rehydration on board, epitope retrieval was performed using Ventana Ultra CC1 buffer (catalog# 6414575001, Roche Diagnostics) at 96°C for 64 minutes. Primary antibody, anti-CD45 (1:200 dilution; catalog# 702575S, Cell Signaling Technologies) was applied at 36°C for 40 minutes. Primary antibody was detected using an anti-rabbit HQ detection system (catalog# 7017936001 and 7017812001, Roche Diagnostics) followed by OPAL 520 (NEL871001KT, Akoya Biosciences) diluted 1:150 in 1X Plus Amplification Diluent (catalog # FP1498, Akoya Biosciences). Following CD45 detection, primary and secondary antibodies from the first round of staining were stripped on board using Ventana Ultra CC1 buffer at 95oC for 12 minutes followed by neutralization using Discovery Inhibitor (catalog #7017944001, Roche Diagnostics). Primary antibody, anti-CD3 (1:200 dilution; catalog# ab16669, Abcam) was applied at 36oC for 40 minutes. CD3 primary antibodies were detected using an anti-rabbit HQ detection system (catalog# 7017936001 and 7017812001, Roche Diagnostics) followed by OPAL 570 (NEL871001KT, Akoya Biosciences) diluted 1:150 in 1X Plus Amplification Diluent (catalog # FP1498, Akoya Biosciences). Primary and secondary antibodies from the second staining round were stripped on board using Ventana Ultra CC1 buffer at 95oC for 12 minutes, then neutralization using Discovery Inhibitor (catalog #7017944001, Roche Diagnostics). Primary antibody, anti-CD11c (1:300 dilution; catalog# PAS-79537, ThermoFisher Scientific) was applied at 36oC for 40 minutes. CD11c primary antibodies were detected using an anti-rabbit HQ detection system (catalog# 7017936001 and 7017812001, Roche Diagnostics) followed by OPAL 690 (NEL871001KT, Akoya Biosciences) diluted 1:150 in 1X Plus Amplification Diluent (catalog # FP1498, Akoya Biosciences). Following CD11c detection, primary and secondary antibodies from the third staining round were stripped on board using Ventana Ultra CC1 buffer at 95oC for 12 minutes and neutralization using Discovery Inhibitor (catalog #7017944001, Roche Diagnostics). Primary antibody, anti-F480 (1:1000 dilution; catalog# 70076, Cell Signaling Technologies) was applied at 36oC for 40 minutes. F4/80 primary antibodies were detected using an anti-rabbit HQ detection system (catalog# 7017936001 and 7017812001, Roche Diagnostics) followed by OPAL Polaris 780 (NEL871001KT, Akoya Biosciences) diluted 1:150 in 1X Plus Amplification Diluent (catalog # FP1498, Akoya Biosciences). Tissue was then counterstained with spectral DAPI (FP1490, Akoya Biosciences) and mounted with Prolong Gold (P36930, ThermoFisher Scientific).

Then, all the slides were scanned using the FL Zeiss whole slide scanner with 40x resolution. Images were initially viewed using Zeiss Zen lite software; HALO® platform v3.4.2986 (Indica Labs, Albuquerque, NM, USA) was used for quantitative analysis using the HighPlex FL algorithm. The real-time proximity analysis/spatial relationship among immune cells was determined based on the 10 μm cut-off, which is considered the regular size of the immune cells, and cell distance equal to or less than that implies cell-cell interaction. Whole slides were scanned and utilized for the quantitative analysis.

### Statistical analysis

All reported P values are from two-sided comparisons. Pairwise comparisons of group mean values for log_10_ CFU (microbiology data) and flow cytometry data were made using one-way analysis of variance followed by Dunnett’s multiple comparisons tests, one-way ANOVA, paired T-test, or Mann-Whitney test where indicated. Several simple linear regressions were performed to assess the correlation of each individual immune response per tissue expressed in percentages (independent variable) with lung mycobacterial outcome (log_10_ transformed-dependent variable). The R^2^ was set above 0.6 to ensure linear correlation and P value was set as <0.05. Prism 10.2.0 (GraphPad Software, Inc. San Diego, CA) was utilized for statistical analyses and figure generation. All error bars represent the estimation of the standard error of the mean, and all midlines represent the group mean. A significance level of α ≤ 0.05 was set for all experiments.

**List of Supplementary Materials:** Fig. S1 to S7

## General

The content of this publication has been available online on the Biorxiv preprint server xx. Illustrations were created with BioRender.com. Illustrations and graphs revised by the Johns Hopkins Illustration Department. We thank Bareng Aletta S. Nonyane, Research Professor in the Global Disease Epidemiology and Control Department of Johns Hopkins School of Public Health, for reviewing the linear regression analysis statistics. We thank Sandeep Tyagi from Nuermberger Lab for gifting us the H37Rv Mtb strain. We thank the JHU research animal resources, veterinary post-doctoral fellows, and the JHU breeding farm staff. Last, we thank Dr. Prakash Srinivasan, the Johns Hopkins Malaria Research Institute, and the Department of Molecular Microbiology and Immunology for Attune™ NxT flow cytometry support.

## Funding

National Institute of Health grant R01AI148710 (PCK, RBM)

National Institute of Health grant K24AI143447 (PCK)

National Institute of Health grant P30AI18436-01 (PCK)

National Institute of Health grant K08AI174959 (SK)

National Institute of Health grant P30CA006973 (Cancer Center Core, William Nelson)

Gilead HIV Research Scholar (SK)

Tuberculosis Research Advancement Center Developmental Award, Johns Hopkins University (SK)

Center for HIV/AIDS Developmental Award, Johns Hopkins University (SK) Clinician Scientist Award, Johns Hopkins University (SK)

Potts Memorial Foundation (SK)

## Author Contributions

Conceptualization: SK, RBM, PCK

Methodology: SK, RBM, PCK, EN, TK, DP, TW, RT, JTG, ARM

Investigation: SK, TK, DP, TW, RT, AY, JRC, JTG, HB, DQ, ARM, KF, HTH, ERS, REB, ADT, YL, JM, HT, JJZ, JZ

Visualization: SK, TW, TK, DP

Funding acquisition: SK, RBM, PCK

Project administration: SK, RBM, PCK

Supervision: SK, RBM, PCK

Writing – original draft: SK

Writing – review & editing: SK, RBM, PCK, EN, TK, DP, TW, AY, JRC, JTG, HB, DQ, RT.

## Data and materials availability

All data are available in the main text or the supplementary materials.

**Fig. S1. Relationships between Rel_Mtb_-specific T-cell responses and dendritic cells in PBMC, BAL, and mediastinal LNs and total CFUs at 12 weeks post- prime vaccination and/or initiation of drug treatment in immunocompetent mice.** Linear regressions were used to test whether antigen-specific CD4+ or CD8+ T-cell produced cytokines (%) are associated with lung mycobacterial burder (total CFUs) in spleen, PBMCs, lungs, mediastinal LNs, and BAL. Results indicate that CD4+ and CD8+ T-cell producing IL17-A, IFN-γ, TNF-α (%) in PBMCs and BAL along with total DCs (%) in BAL and CD4+ and CD8+ T-cell producing IL17-A and TNF-α (%) are significantly negatively associated with lung mycobacterial burden. R^2^ is the coefficient of determination. Each dot represents the pooled sample (single-cell suspension) from 2-7 animals. Black lines represent linear fit (with 95% confidence interval in dotted black lines). PBMC, peripheral blood mononuclear cells; BAL, bronchoalveolar lavage; LN, lymph nodes; CFU, colony-forming units.

**Fig. S2. T cells in mouse lungs did not statistically differ between RHZE vs. IN *MIP-3*α*/rel_Mtb_* + RHZE after 12 weeks of treatment in immunocompetent mice as measured through immunofluorescence.** T cells are defined as DAPI+CD45+CD3+ and quantified as T cells/ DAPI+ cells. [RHZE (n=6) vs. IN MIP-3α/rel_Mtb_ & RHZE (n=4), Mann-Whitney test was run on medians, the assessment was performed in whole slides]; RHZE, Rifampin-Isoniazid- Pyrazinamide-Ethambutol.

**Fig. S3. Spleen mycobacterial burden was more effectively reduced in CD4 KO mice treated with IN *MIP-3*α*/rel_Mtb_* fusion vaccine + RHZE compared to RHZE alone 6 weeks after treatment initiation.** IN, intranasal; CFU, colony- forming units; RHZE, Rifampin-Isoniazid-Pyrazinamide-Ethambutol; CD4 KO, CD4+ T cell knockout; ****P<0.0001.

**Fig. S4. Spleen mycobacterial burden did not statistically differ in immunocompetent mice treated with BPaL alone vs. IN *MIP-3*α*/rel_Mtb_* fusion vaccine + BPaL.** IN, intranasal; CFU, colony-forming units; BPaL, Bedaquiline- Pretomanid-Linezolid; ****P<0.0001.

**Fig. S5. IN *MIP-3*α*/rel_Mtb_* DNA fusion vaccine induced sustained TB-protective immune responses in nonhuman primates by 24 weeks after primary vaccination.** Pre-vaccination, 9-week and 24-week rel_Mtb-_specific CD4+ and CD8+T-cell responses (IFN-γ, TNF-α, IL-17A) in **(A)** PBMC and **(B)** BAL did not significantly defer over time. Cytokines are expressed in percentages. IN, intranasal; PBMC, peripheral blood mononuclear cells; BAL, bronchoalveolar lavage; prevax, prevaccination.

**Fig. S6. Two-dimensional gating strategy for mouse flow cytometric identification of (A) T-cell-producing cytokines, (B) DCs, (C) activated DCs, and (D) lymphoid or non-lymphoid tissue cDC1 and cDC2.** We excluded doublets and debris and gated on single live CD45+ cells. **(A)** We identified T cells (CD45+CD3+), CD4+ T cells, CD8+ T cells, and T-cell producing cytokines (CD45+CD3+CD4+ IFN-γ+, TNF-α+, or IL-17A+ and CD45+CD3+CD8+ IFN-γ+, TNF-α+, or IL-17A+); **(B)** DCs (CD45+CD3-CD19-NK1.1-Ly6G- MHCII+CD11c+CD24+ or CD45+CD3-CD19-NK1.1-Ly6G-MHCII+CD11c+CD64-) **(C)** activated DCs (CD45+CD3-CD19-NK1.1-Ly6G- MHCII+CD11c+CD24+CD80+ or CD45+CD3-CD19-NK1.1-Ly6G-MHCII+CD11c+CD64-CD80+) and **(D)** non-lymphoid tissue (lungs and BAL) DC I (CD45+CD3-CD19-NK1.1-Ly6G-MHCII+CD11c+CD24+CD103+CD11b-) and DCII (CD45+CD3-CD19-NK1.1-Ly6G-MHCII+CD11c+CD24+CD103-CD11b+), and lymphoid tissue (spleen, LNs, PBMCs) DC I (CD45+CD3-CD19-NK1.1- Ly6G-MHCII+CD11c+CD24+XCR1+CD11b-) and DC II (CD45+CD3-CD19-NK1.1-Ly6G-MHCII+CD11c+CD24+XCR1-CD11b+); DCs, total classical dendritic cells; cDCs I, classical dendritic cells type I; cDCs II, classical dendritic cell type II; PBMC, peripheral blood mononuclear cells; BAL, bronchoalveolar lavage; LNs, mediastinal LNs

**Fig. S7. Two-dimensional gating strategy for nonhuman primate flow cytometric identification of T-cell-producing cytokines.** We excluded doublets and debris and gated on single live cells. We identified T cells (CD3+), CD4+ T cells, CD8+ T cells and T-cell producing cytokines (CD3+CD4+IFN-γ+, TNF-α+, or IL-17A+ and CD3+CD8+IFN-γ+, TNF-α+, or IL-17A+).

## References

1. WHO. (2023), vol. 2023.

2. A. S. Fauci, Multidrug-resistant and extensively drug-resistant tuberculosis: the National Institute of Allergy and Infectious Diseases Research agenda and recommendations for priority research. The Journal of infectious diseases 197, 1493–1498 (2008).

3. Y. M. Chuang, N. K. Dutta, J. T. Gordy, V. L. Campodónico, M. L. Pinn, R. B. Markham, C. F. Hung, P. C. Karakousis, Antibiotic Treatment Shapes the Antigenic Environment During Chronic TB Infection, Offering Novel Targets for Therapeutic Vaccination. Frontiers in immunology 11, 680 (2020).

4. A. H. Diacon, A. Pym, M. P. Grobusch, J. M. de los Rios, E. Gotuzzo, I. Vasilyeva, V. Leimane, K. Andries, N. Bakare, T. De Marez, M. Haxaire- Theeuwes, N. Lounis, P. Meyvisch, E. De Paepe, R. P. van Heeswijk, B. Dannemann, Multidrug-resistant tuberculosis and culture conversion with bedaquiline. N Engl J Med 371, 723–732 (2014).

5. A. S. Pym, A. H. Diacon, S. J. Tang, F. Conradie, M. Danilovits, C. Chuchottaworn, I. Vasilyeva, K. Andries, N. Bakare, T. De Marez, M. Haxaire- Theeuwes, N. Lounis, P. Meyvisch, B. Van Baelen, R. P. van Heeswijk, B. Dannemann, Bedaquiline in the treatment of multidrug- and extensively drug-resistant tuberculosis. Eur Respir J 47, 564–574 (2016).

6. T. V. A. Nguyen, R. M. Anthony, A.-L. Bañuls, T. V. A. Nguyen, D. H. Vu, J.-W. C. Alffenaar, Bedaquiline Resistance: Its Emergence, Mechanism, and Prevention. Clinical Infectious Diseases 66, 1625–1630 (2018).

7. W. H. Organization. (2019), vol. 2024.

8. E. Guirado, O. Gil, N. Cáceres, M. Singh, C. Vilaplana, P. J. Cardona, Induction of a specific strong polyantigenic cellular immune response after short-term chemotherapy controls bacillary reactivation in murine and guinea pig experimental models of tuberculosis. Clin Vaccine Immunol 15, 1229–1237 (2008).

9. A. Gupta, F. J. Ahmad, F. Ahmad, U. D. Gupta, M. Natarajan, V. Katoch, S. Bhaskar, Efficacy of Mycobacterium indicus pranii immunotherapy as an adjunct to chemotherapy for tuberculosis and underlying immune responses in the lung. PLoS One 7, e39215 (2012).

10. W. P. Gong, Y. Liang, Y. B. Ling, J. X. Zhang, Y. R. Yang, L. Wang, J. Wang, Y. C. Shi, X. Q. Wu, Effects of Mycobacterium vaccae vaccine in a mouse model of tuberculosis: protective action and differentially expressed genes. Mil Med Res 7, 25 (2020).

11. R. N. Coler, S. Bertholet, S. O. Pine, M. T. Orr, V. Reese, H. P. Windish, C. Davis, M. Kahn, S. L. Baldwin, S. G. Reed, Therapeutic immunization against Mycobacterium tuberculosis is an effective adjunct to antibiotic treatment. The Journal of infectious diseases 207, 1242–1252 (2013).

12. S. Karanika, J. T. Gordy, P. Neupane, T. Karantanos, J. Ruelas Castillo, D. Quijada, K. Comstock, A. K. Sandhu, A. R. Kapoor, Y. Hui, S. K. Ayeh, R. Tasneen, S. Krug, C. Danchik, T. Wang, C. Schill, R. B. Markham, P. C. Karakousis, An intranasal stringent response vaccine targeting dendritic cells as a novel adjunctive therapy against tuberculosis. Frontiers in immunology 13, (2022).

13. C. Danchik, S. Wang, P. C. Karakousis, Targeting the Mycobacterium tuberculosis Stringent Response as a Strategy for Shortening Tuberculosis Treatment. Frontiers in microbiology 12, 744167 (2021).

14. J. Prusa, D. X. Zhu, C. L. Stallings, The stringent response and Mycobacterium tuberculosis pathogenesis. Pathogens and disease 76, (2018).

15. B. B. S. Passos, M. Araújo-Pereira, C. L. Vinhaes, E. P. Amaral, B. B. Andrade, The role of ESAT-6 in tuberculosis immunopathology. Frontiers in immunology 15, (2024).

16. A. M. Cooper, Cell-mediated immune responses in tuberculosis. Annu Rev Immunol 27, 393–422 (2009).

17. S. A. Khader, G. K. Bell, J. E. Pearl, J. J. Fountain, J. Rangel-Moreno, G. E. Cilley, F. Shen, S. M. Eaton, S. L. Gaffen, S. L. Swain, R. M. Locksley, L. Haynes, T. D. Randall, A. M. Cooper, IL-23 and IL-17 in the establishment of protective pulmonary CD4+ T cell responses after vaccination and during Mycobacterium tuberculosis challenge. Nat Immunol 8, 369–377 (2007).

18. H. P. Gideon, J. Phuah, A. J. Myers, B. D. Bryson, M. A. Rodgers, M. T. Coleman, P. Maiello, T. Rutledge, S. Marino, S. M. Fortune, D. E. Kirschner, P. L. Lin, J. L. Flynn, Variability in tuberculosis granuloma T cell responses exists, but a balance of pro- and anti-inflammatory cytokines is associated with sterilization. PLoS Pathog 11, e1004603 (2015).

19. D. A. Anderson, 3rd, K. M. Murphy, C. G. Briseño, Development, Diversity, and Function of Dendritic Cells in Mouse and Human. Cold Spring Harb Perspect Biol 10, (2018).

20. N. van der Wel, D. Hava, D. Houben, D. Fluitsma, M. van Zon, J. Pierson, M. Brenner, P. J. Peters, M. tuberculosis and M. leprae translocate from the phagolysosome to the cytosol in myeloid cells. Cell 129, 1287–1298 (2007).

21. F. Winau, S. Weber, S. Sad, J. de Diego, S. L. Hoops, B. Breiden, K. Sandhoff, V. Brinkmann, S. H. Kaufmann, U. E. Schaible, Apoptotic vesicles crossprime CD8 T cells and protect against tuberculosis. Immunity 24, 105–117 (2006).

22. F. Conradie, R. Bagdasaryan Tatevik, S. Borisov, P. Howell, L. Mikiashvili, N. Ngubane, A. Samoilova, S. Skornykova, E. Tudor, E. Variava, P. Yablonskiy, D. Everitt, H. Wills Genevieve, E. Sun, M. Olugbosi, E. Egizi, M. Li, A. Holsta, J. Timm, A. Bateson, M. Crook Angela, M. Fabiane Stella, R. Hunt, D. McHugh Timothy, D. Tweed Conor, S. Foraida, M. Mendel Carl, M. Spigelman, Bedaquiline–Pretomanid–Linezolid Regimens for Drug-Resistant Tuberculosis. New England Journal of Medicine 387, 810–823 (2022).

23. WHO. (2023), vol. 2024.

24. R. Lai, A. F. Ogunsola, T. Rakib, S. M. Behar, Key advances in vaccine development for tuberculosis—success and challenges. npj Vaccines 8, 158 (2023).

25. I. Messaoudi, R. Estep, B. Robinson, S. W. Wong, Nonhuman primate models of human immunology. Antioxid Redox Signal 14, 261–273 (2011).

26. K. A. Phillips, K. L. Bales, J. P. Capitanio, A. Conley, P. W. Czoty, B. A. t Hart, W. D. Hopkins, S. L. Hu, L. A. Miller, M. A. Nader, P. W. Nathanielsz, J. Rogers, C. A. Shively, M. L. Voytko, Why primate models matter. Am J Primatol 76, 801–827 (2014).

27. D. J. Dowling, O. Levy, Ontogeny of early life immunity. Trends Immunol 35, 299–310 (2014).

28. A. W. Stowers, V. Cioce, R. L. Shimp, M. Lawson, G. Hui, O. Muratova, D. C. Kaslow, R. Robinson, C. A. Long, L. H. Miller, Efficacy of two alternate vaccines based on Plasmodium falciparum merozoite surface protein 1 in an Aotus challenge trial. Infection and immunity 69, 1536–1546 (2001).

29. J. E. Rosas, J. L. Pedraz, R. M. Hernandez, A. R. Gascon, M. Igartua, F. Guzman, R. Rodriguez, J. Cortes, M. E. Patarroyo, Remarkably high antibody levels and protection against P. falciparum malaria in Aotus monkeys after a single immunisation of SPf66 encapsulated in PLGA microspheres. Vaccine 20, 1707–1710 (2002).

30. R. Abrahem, E. Chiang, J. Haquang, A. Nham, Y. S. Ting, V. Venketaraman, The Role of Dendritic Cells in TB and HIV Infection. J Clin Med 9, (2020).

31. K. Uehira, R. Amakawa, T. Ito, K. Tajima, S. Naitoh, Y. Ozaki, T. Shimizu, K. Yamaguchi, Y. Uemura, H. Kitajima, S. Yonezu, S. Fukuhara, Dendritic cells are decreased in blood and accumulated in granuloma in tuberculosis. Clin Immunol 105, 296–303 (2002).

32. M. Lichtner, R. Rossi, F. Mengoni, S. Vignoli, B. Colacchia, A. P. Massetti, I. Kamga, A. Hosmalin, V. Vullo, C. M. Mastroianni, Circulating dendritic cells and interferon-alpha production in patients with tuberculosis: correlation with clinical outcome and treatment response. Clin Exp Immunol 143, 329–337 (2006).

33. A. Biragyn, M. Surenhu, D. Yang, P. A. Ruffini, B. A. Haines, E. Klyushnenkova, J. J. Oppenheim, L. W. Kwak, Mediators of innate immunity that target immature, but not mature, dendritic cells induce antitumor immunity when genetically fused with nonimmunogenic tumor antigens. J Immunol 167, 6644–6653 (2001).

34. P. A. Darrah, J. J. Zeppa, P. Maiello, J. A. Hackney, M. H. Wadsworth, T. K. Hughes, S. Pokkali, P. A. Swanson, N. L. Grant, M. A. Rodgers, M. Kamath, C. M. Causgrove, D. J. Laddy, A. Bonavia, D. Casimiro, P. L. Lin, E. Klein, A. G. White, C. A. Scanga, A. K. Shalek, M. Roederer, J. L. Flynn, R. A. Seder, Prevention of tuberculosis in macaques after intravenous BCG immunization. Nature 577, 95–102 (2020).

35. J. Wang, L. Thorson, R. W. Stokes, M. Santosuosso, K. Huygen, A. Zganiacz, M. Hitt, Z. Xing, Single mucosal, but not parenteral, immunization with recombinant adenoviral-based vaccine provides potent protection from pulmonary tuberculosis. J Immunol 173, 6357–6365 (2004).

36. M. Jeyanathan, A. Heriazon, Z. Xing, Airway luminal T cells: a newcomer on the stage of TB vaccination strategies. Trends Immunol 31, 247–252 (2010).

37. N. F. Walker, G. Meintjes, R. J. Wilkinson, HIV-1 and the immune response to TB. Future Virol 8, 57–80 (2013).

38. K. Loré, A. Sönnerborg, C. Broström, L. E. Goh, L. Perrin, H. McDade, H. J. Stellbrink, B. Gazzard, R. Weber, L. A. Napolitano, Y. van Kooyk, J. Andersson, Accumulation of DC-SIGN+CD40+ dendritic cells with reduced CD80 and CD86 expression in lymphoid tissue during acute HIV-1 infection. Aids 16, 683–692 (2002).

39. J.-C. Cancel, K. Crozat, M. Dalod, R. Mattiuz, Are Conventional Type 1 Dendritic Cells Critical for Protective Antitumor Immunity and How? Frontiers in immunology 10, (2019).

40. J. Millard, S. Rimmer, C. Nimmo, M. O’Donnell, Therapeutic Failure and Acquired Bedaquiline and Delamanid Resistance in Treatment of Drug-Resistant TB. Emerg Infect Dis 29, 1081–1084 (2023).

41. S. V. Omar, F. Ismail, N. Ndjeka, K. Kaniga, N. A. Ismail, Bedaquiline-Resistant Tuberculosis Associated with Rv0678 Mutations. N Engl J Med 386, 93–94 (2022).

42. D. Hobernik, M. Bros, DNA Vaccines-How Far From Clinical Use? Int J Mol Sci 19, (2018).

43. K. A. Kraynyak, E. Blackwood, J. Agnes, P. Tebas, M. Giffear, D. Amante, E. L. Reuschel, M. Purwar, A. Christensen-Quick, N. Liu, V. M. Andrade, M. C. Diehl, S. Wani, M. Lupicka, A. Sylvester, M. P. Morrow, P. Pezzoli, T. McMullan, A. J. Kulkarni, F. I. Zaidi, D. Frase, K. Liaw, T. R. F. Smith, S. J. Ramos, J. Ervin, M. Adams, J. Lee, M. Dallas, A. Shah Brown, J. E. Shea, J. J. Kim, D. B. Weiner, K. E. Broderick, L. M. Humeau, J. D. Boyer, M. P. Mammen, SARS-CoV-2 DNA Vaccine INO-4800 Induces Durable Immune Responses Capable of Being Boosted in a Phase 1 Open-Label Trial. The Journal of infectious diseases 225, 1923–1932 (2022).

44. A. Khobragade, S. Bhate, V. Ramaiah, S. Deshpande, K. Giri, H. Phophle, P. Supe, I. Godara, R. Revanna, R. Nagarkar, J. Sanmukhani, A. Dey, T. M. C. Rajanathan, K. Kansagra, P. Koradia, Efficacy, safety, and immunogenicity of the DNA SARS-CoV-2 vaccine (ZyCoV-D): the interim efficacy results of a phase 3, randomised, double-blind, placebo-controlled study in India. Lancet 399, 1313–1321 (2022).

45. J. Y. Ahn, J. Lee, Y. S. Suh, Y. G. Song, Y.-J. Choi, K. H. Lee, S. H. Seo, M. Song, J.-W. Oh, M. Kim, H. Y. Seo, J.-E. Kwak, J. W. Youn, J. W. Woo, E.-C. Shin, Y. C. Sung, S.-H. Park, J. Y. Choi, Safety and immunogenicity of two recombinant DNA COVID-19 vaccines containing the coding regions of the spike or spike and nucleocapsid proteins: an interim analysis of two open-label, non-randomised, phase 1 trials in healthy adults. The Lancet Microbe 3, e173–e183 (2022).

46. S. Kodama, N. Abe, T. Hirano, M. Suzuki, A single nasal dose of CCL20 chemokine induces dendritic cell recruitment and enhances nontypable Haemophilus influenzae-specific immune responses in the nasal mucosa. Acta Otolaryngol 131, 989–996 (2011).

47. A. A. Ordonez, R. Tasneen, S. Pokkali, Z. Xu, P. J. Converse, M. H. Klunk, D. J. Mollura, E. L. Nuermberger, S. K. Jain, Mouse model of pulmonary cavitary tuberculosis and expression of matrix metalloproteinase-9. Dis Model Mech 9, 779–788 (2016).

48. M. E. Urbanowski, A. A. Ordonez, C. A. Ruiz-Bedoya, S. K. Jain, W. R. Bishai, Cavitary tuberculosis: the gateway of disease transmission. Lancet Infect Dis 20, e117–e128 (2020).

49. K. N. Williams, S. J. Brickner, C. K. Stover, T. Zhu, A. Ogden, R. Tasneen, S. Tyagi, J. H. Grosset, E. L. Nuermberger, Addition of PNU-100480 to first-line drugs shortens the time needed to cure murine tuberculosis. Am J Respir Crit Care Med 180, 371–376 (2009).

50. D. Almeida, J. Converse Paul, S.-Y. Li, M. Upton Anna, N. Fotouhi, L. Nuermberger Eric, Comparative Efficacy of the Novel Diarylquinoline TBAJ-876 and Bedaquiline against a Resistant Rv0678 Mutant in a Mouse Model of Tuberculosis. Antimicrobial Agents and Chemotherapy 65, 10.1128/aac.01412-01421 (2021).

51. S. Tyagi, E. Nuermberger, T. Yoshimatsu, K. Williams, I. Rosenthal, N. Lounis, W. Bishai, J. Grosset, Bactericidal Activity of the Nitroimidazopyran PA-824 in a Murine Model of Tuberculosis. Antimicrobial Agents and Chemotherapy 49, 2289–2293 (2005).

52. F. Conradie, T. R. Bagdasaryan, S. Borisov, P. Howell, L. Mikiashvili, N. Ngubane, A. Samoilova, S. Skornykova, E. Tudor, E. Variava, P. Yablonskiy, D. Everitt, G. H. Wills, E. Sun, M. Olugbosi, E. Egizi, M. Li, A. Holsta, J. Timm, A. Bateson, A. M. Crook, S. M. Fabiane, R. Hunt, T. D. McHugh, C. D. Tweed, S. Foraida, C. M. Mendel, M. Spigelman, Bedaquiline–Pretomanid–Linezolid Regimens for Drug-Resistant Tuberculosis. New England Journal of Medicine 387, 810–823 (2022).

53. L. Van Hoecke, E. R. Job, X. Saelens, K. Roose, Bronchoalveolar Lavage of Murine Lungs to Analyze Inflammatory Cell Infiltration. J Vis Exp, (2017).

