## Supplementary figures and images for "Therapeutic DNA Vaccine Targeting *Mycobacterium tuberculosis* Persisters Shortens Curative Tuberculosis Treatment"

### Fig. S1

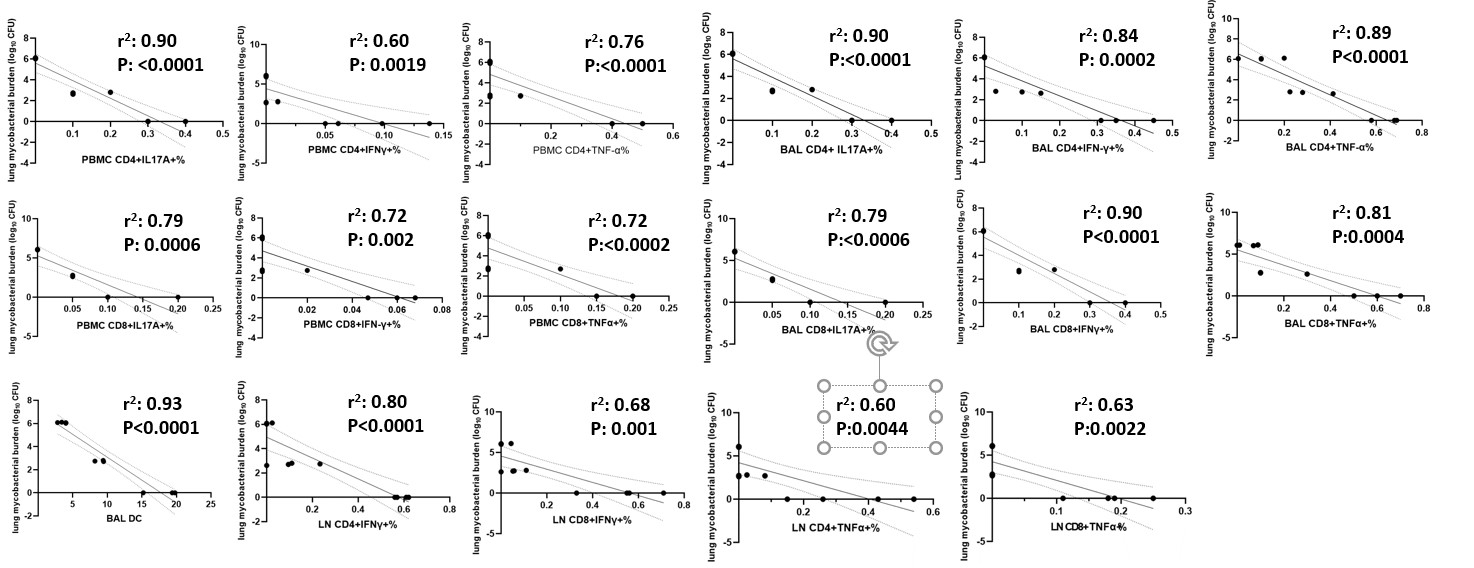

### Fig. S2

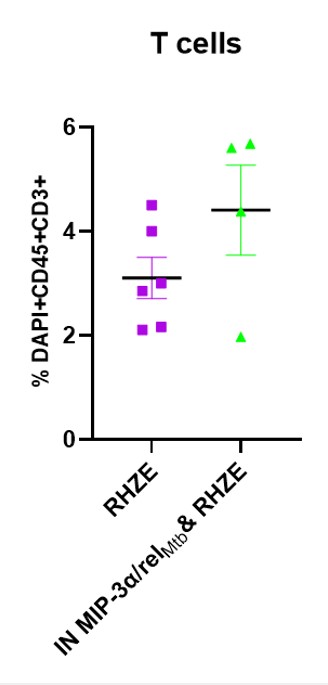

### Fig. S5

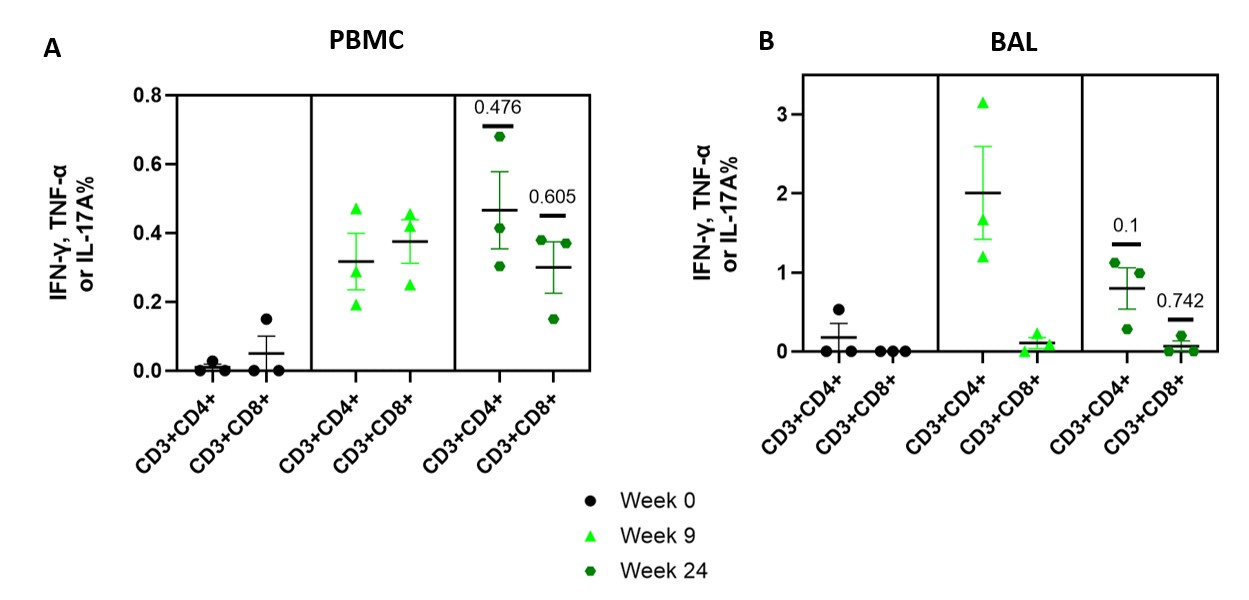

### Fig. S6

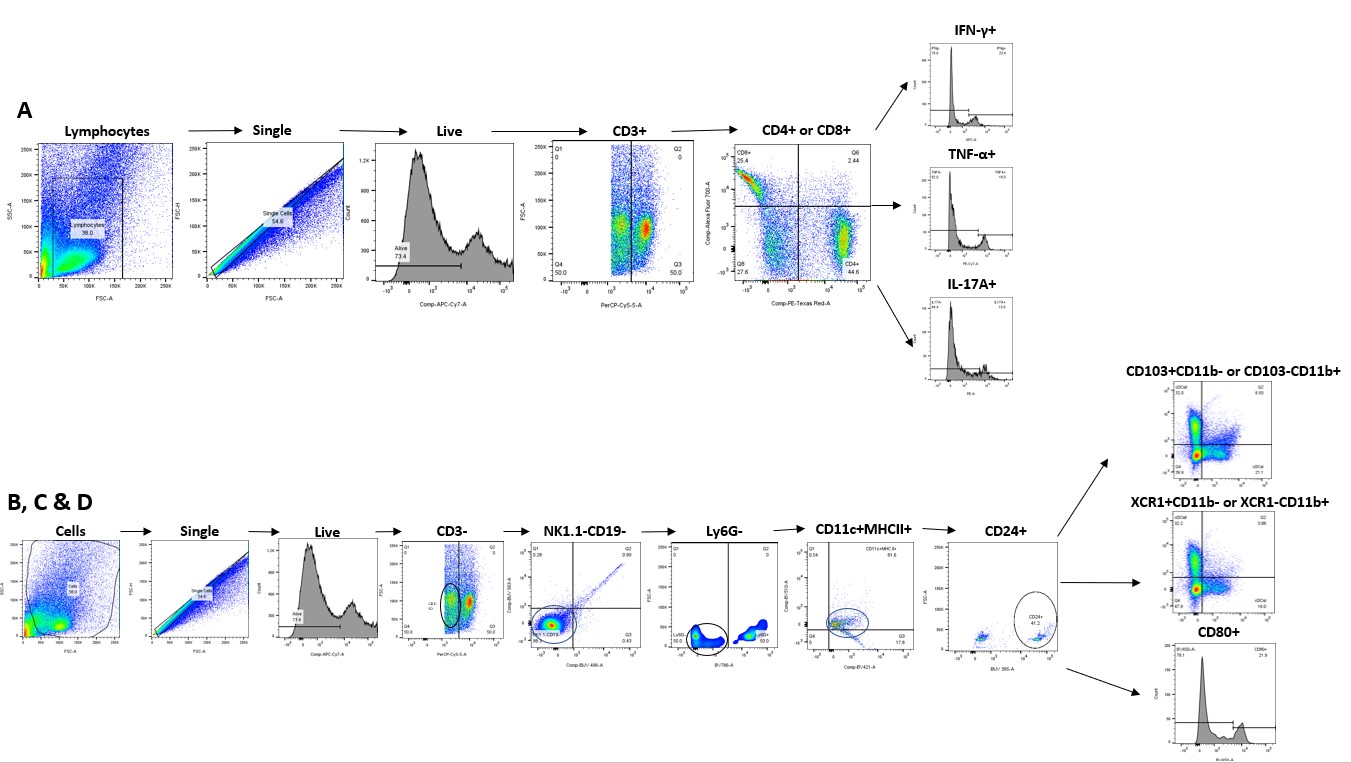

### Fig. S7

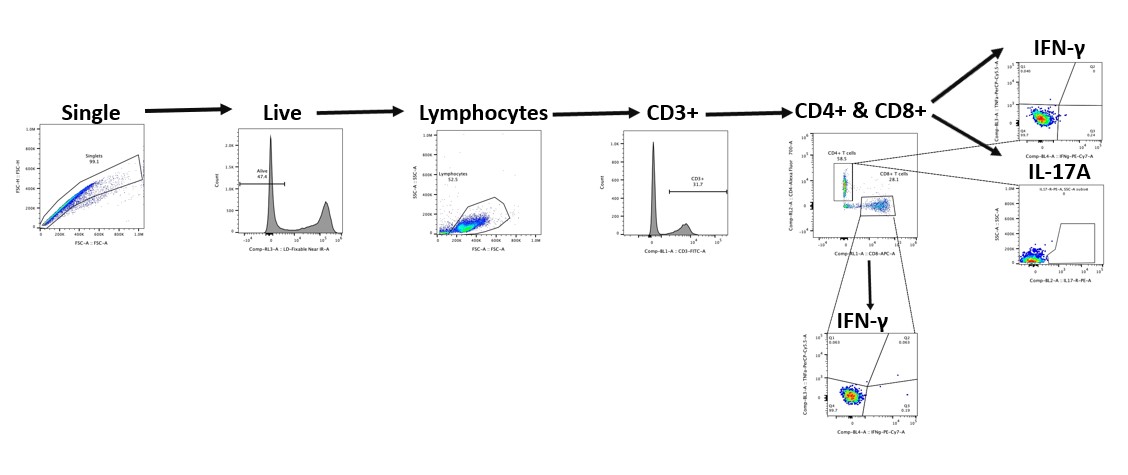

### Fig.S3

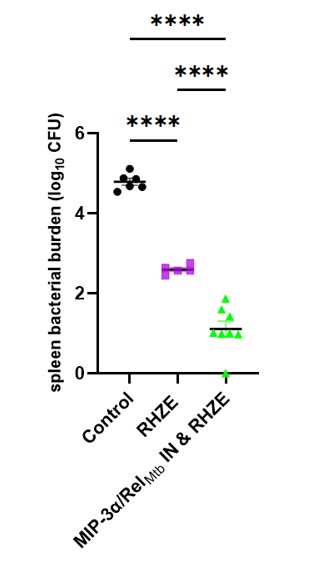

### Fig.S4

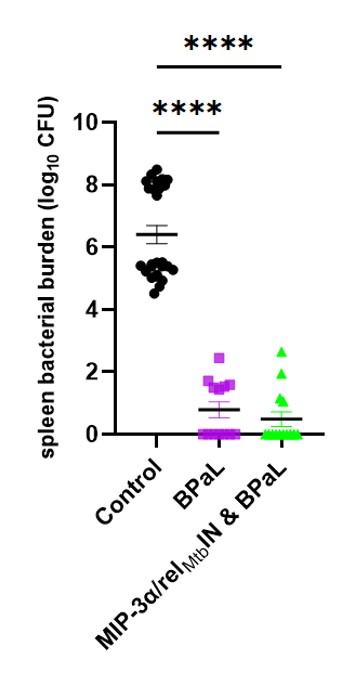
